# Alternative stable states in a model of microbial community limited by multiple essential nutrients

**DOI:** 10.1101/439547

**Authors:** Veronika Dubinkina, Yulia Fridman, Parth Pratim Pandey, Sergei Maslov

**Author notes:** These three authors contributed equally (Veronika Dubinkina, Yulia Fridman, Parth Pratim Pandey).

## Abstract

Microbial communities routinely have several alternative stable states observed for the same environmental parameters. Sudden and irreversible transitions between these states make external manipulation of these systems more complicated. To better understand the mechanisms and origins of multistability in microbial communities, we introduce and study a model of a microbial ecosystem colonized by multiple specialist species selected from a fixed pool. Growth of each species can be limited by essential nutrients of two types, e.g. carbon and nitrogen, each represented in the environment by multiple metabolites. We demonstrate that our model has an exponentially large number of potential stable states realized for different environmental parameters. Using game theoretical methods adapted from the stable marriage problem we predict all of these states based only on ranked lists of competitive abilities of species for each of the nutrients. We show that for every set of nutrient influxes, several mutually uninvadable stable states are generally feasible and we distinguish them based upon their dynamic stability. We further explore an intricate network of discontinuous transitions (regime shifts) between these alternative states both in the course of community assembly, or upon changes of nutrient influxes.

## INTRODUCTION

Microbial communities play an important role in medicine (human microbiome^1, 2^), agriculture (soil^3^, plant root^4^, and animal^5^ microbiomes), climate, (via carbon cycle feedbacks^6^), and technology (industrial bioreactors^7^, wastewater digesters^8^, etc.) They are often characterized by more than one stable state observed for the same set of environmental parameters^9–12^. Such alternative stable states^13–17^ have several hallmark properties discussed in Ref.^18^ including “discontinuity in the response to an environmental driving parameter” (referred to as regime shifts in the ecosystems literature), lack of recovery after a perturbation (hysteresis), and “divergence due to different initial conditions” or due to the order in which species were introduced during the initial colonization process^19^.

To be able to predict the behavior of a microbial community, one needs to understand the mechanisms that favor one such state over the other and the factors triggering transitions between them. In many practical situations we would also like to be able to manipulate and control a microbial ecosystem in a predictable manner, and the existence of more than one stable state greatly complicates this task^20^.

Growth rates and, ultimately, abundances of microbial species are affected by multiple factors, with availability of nutrients being among the most important ones. Thus environmental concentrations and influx rates of externally supplied nutrients play a crucial role in determining the state (or multiple states) of a microbial ecosystem defined by its species composition. Changes in nutrient concentrations can also trigger transitions (regime shifts^17^) between these states (see Ref.^21^ for a recent literature survey on microbial communities’ response to disturbances). Nutrients required for growth of a microbial (or any other) species exist in the form of multiple metabolites of several essential types (i.e. sources of C, N, P, Fe, etc.). The growth of each species is usually limited by the most scarce type of nutrient (for an exception to this rule see Ref.^22^ demonstrating that oceanic phytoplankton can be co-limited by more than one essential nutrient).

Here we introduce and study a new mathematical model of a microbial community limited by multiple essential nutrients. To put our model in context of previously studied ones, we briefly review common approaches to modelling of microbial communities.

One of the simplest and thereby most popular approaches in ecological modelling^23–36^ is based on variants of generalized Lotka-Volterra (gLV) model^37, 38^. The gLV model does not explicitly consider nutrients, replacing them with the effective direct inter-species interactions While gLV models have provided valuable insights due to their simplicity, they have also been criticized when applied to multispecies communities^39^.

Another popular approach is based on variants of the classic MacArthur consumer-resource model^40, 41^ in which the growth rate of each species is given by a linear combination of concentrations of several fully substitutable resources^19, 42–46^. This corresponds to a logical OR-gate operating on all nutrient inputs of a given species. A more general case in which growth rate can be arbitrary non-linear function of concentrations of *just two resources* has been considered in the foundational work by Tilman^47^. The modeling framework and the geometric interpretation of resource dynamics developed in Ref.^47^ proved to be useful for interpreting experimental data describing ecology and plankton communities^48^ and remains an active field of research^49–51^. While bistability between a pair of species has already been mentioned in Ref.^47^, more complex scenarios with multiple species and/or more than two alternative stable states, to the best of our knowledge has not been described (see^35^ for the analysis of alternative stable states in gLV model).

Our study fills this gap by generalizing the model of Ref.^47^ to more than two metabolites. The population dynamics in our model is shaped by species competing for *multiple metabolites* of two *essential types* to which we refer to as sources of carbon and nitrogen, while any other pair of essential nutrients is equally possible. In our model multiple different metabolites (e.g. different sugars) can serve as carbon sources, and another set of metabolites - as nitrogen sources (for simplicity we ignore the possibility of the same metabolite providing both carbon and nitrogen). The ecosystem in our model is colonized by highly specialized species, with each species capable of utilizing just one specific pair of metabolites as its carbon and one nitrogen sources. Using specialist species greatly simplifies our calculations but we will also propose variants of our model incorporating generalist species.

We show that our model is characterized by exponentially large number of steady states, each realized for different sets of environmental parameters. Using game theoretical methods adapted from the well-known stable marriage problem^52, 53^, we show that all of these states can be identified based only on ranked lists of competitive abilities of species for each of the resources. For any set of nutrient influxes a few *mutually uninvadable stable states* are generally feasible. They may or may not be dynamically stable, and our methods allow us to infer dynamic stability of each of them for any set of species’ C:N stoichiometries. As in Ref.^41, 47^, multistability (alternative stable states) is only possible when stoichiometric ratios of different species are not identical. Our model allows us to explore the intricate network of discontinuous transitions (regime shifts) between these alternative states in the course of community assembly and changing nutrient influxes. The aim of our study is to provide an intuitive understanding of the basic rules governing the existence of alternative communities in microbial ecosystems growing on multiple essential resources and of transitions between these states.

While we formulate our model for microbial ecosystems, nothing in its rules prevent it from describing macroscopic ecosystems, e.g. that dominated by plants. In fact, the model of Ref.^47^, which our model generalizes, has been successfully applied to a broad variety of natural and artificial ecosystems.

## MODEL AND RESULTS

Our model describes an ecosystem colonized by microbes selected from a pool of *S* species. Growth of species in our community is limited by two types of essential nutrients, which we will refer to as “carbon” and “nitrogen” sources. In principle, these could be any two types of nutrients essential for life: C, N, P, Fe, etc. A straightforward generalization of our model involves three or more types of essential nutrients. Carbon and nitrogen sources exist in the environment in the form of K distinct metabolites containing carbon, and *M* other metabolites containing nitrogen. To allow for a mathematical understanding of steady states in our model we assume that each of our *S* species is an extreme specialist, capable of utilizing a single pair resources, i.e., one carbon and one nitrogen metabolites. We further assume that the growth rate *g*_*α*_ of a species *α* is determined by the concentration of the rate-limiting resource via Liebig’s law of the minimum^54^:

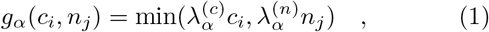

where *c*_*i*_ and *n*_*j*_, are the environmental concentrations of the carbon resource *i* and the nitrogen resource *j* consumed by this species *α*, while 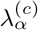 and 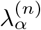 are, respectively, its competitive abilities for these resources. The dynamics of microbial populations *B*_*α*_ is defined by:

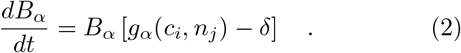

Here we assumed that microbes have no maintenance costs and that both microbes and their resources are in a chemostat-like environment subject to a constant dilution rate *δ*. However, all our results remain unchanged in a more general case of non-zero (and microbe-specific) microbial maintenance cost that could be different from the dilution rate of resources. The resources are externally supplied to our system at fixed influxes 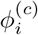 and 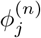 and their concentrations follow the equations:

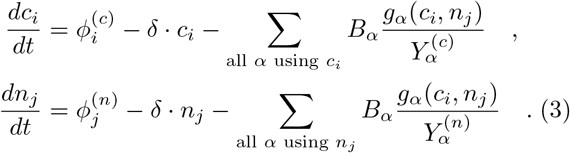

Here 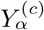 and 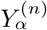 are the carbon and nitrogen growth yields of the species *α* quantifying the number of microbial cells generated per unit of concentration of each of the two consumed resources. It is easy to show that our system satisfies mass conservation laws:

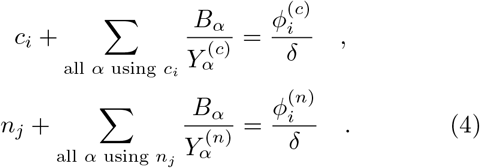

The concentrations of all surviving bacteria and all resources in a steady state are determined by setting the left hand sides of Eqs. 2,3 to zero and solving them for the steady state concentrations *B*_*α*_, *c*_*i*_, and *n*_*j*_. As we will show the steady state equations impose a number of constraints on competitive abilities of surviving microbes, their yields, and nutrient fluxes, where a given steady state is feasible. One type of constraints comes from the competitive exclusion principle^55, 56^ contained in the equations 2. In models with substitutable resources of one type (say, multiple sources of carbon), the specialist species with the largest competitive ability λ^(*c*)^ generally wins the battle for each carbon source (see e.g. Ref.^57^, where it was employed to describe the colonization dynamics of an ecosystem with cross-feeding). In the case of non-substitutable essential resources of two (or more) types, that is the subject of this study, this simple rule is replaced with the following *two competitive exclusion rules*:

- Exclusion Rule 1: Each nutrient (either carbon or nitrogen source) can limit the growth of no more than one species α. From this it follows that (barring special circumstances) the number of surviving species in any given steady state cannot be larger than *K* + *M*, the total number of nutrients.
- Exclusion Rule 2: Each nutrient (say, a specific carbon source) can be used by any number of species in a non-rate-limiting fashion (that is to say, where it does not influence species’ growth rate by setting the value of the minimum in Eq. 1). However, any such species *β* has to have 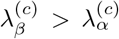 where 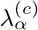 is the competitive ability of the species whose growth is limited by this nutrient. In case of a non-rate-limiting use of a nitrogen source, the constraint becomes 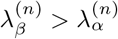.

As we will show below, any set of microbes with the rate-limiting nutrient specified for each microbe, that satisfy the above two constraints imposed by the exclusion rules is a steady state in our system for some set of nutrients influxes (but this does not characterize the dynamic stability of the states). We will refer to them as states allowed by the exclusion rules, or simply “allowed states”.

Each allowed state could be conveniently visualized in terms of a bipartite directed network with vertices corresponding to individual resources and edges connecting carbon and nitrogen sources - to surviving species (see Fig. 1A). We choose the direction of each edge to go from the rate-limiting resource for this species to the non-rate-limiting one. Our exclusion rules can be reformulated in the network language as follows: each vertex can have at most one outgoing link and any number of incoming links (Rule 1). All incoming links have to have larger values of λ than the outgoing link (if any) (Rule 2). Hence, the task of discovering and enumerating all possible steady states realized for different nutrient influxes is equivalent to finding the set of all directed graphs satisfying the above constraints. An allowed state can also be conveniently represented as a matrix with K rows (representing the K carbon nutrients) and M columns (representing the M nitrogen nutrients) where each element (*i*, *j*) represents the specialist species consuming *i*^th^ carbon source and *j*^th^ nitrogen source. To convey the limiting nutrient for the (*i*, *j*)^th^ species we color the (*i*, *j*)^th^ cell of the state matrix by red if it is carbon limited or blue if the species is nitrogen limited. A cell is left empty/uncolored if the species allowed to consume that pair of resources is absent from the community. In this formulation the constraints imposed by the exclusion rules translate to: Rule 1 - each row of the state matrix can have at most one red species (i.e., limited by carbon) and each column can have at most one blue species (i.e., limited by nitrogen) and Rule 2 - In each row λ^(*c*)^ of all blue species should be larger than the λ^(*c*)^ of the red species (if any) and similarly in every column all red species should have a larger λ^(*n*)^ than the λ^(*n*)^ of the blue species (if any). 1B) shows the corresponding matrix form of the state described in Fig 1A. We will use this matrix representation of states in the following figures.

It is useful to single out a subset of “uninvadable states” among all of the steady states allowed by the exclusion rules. These are defined by the condition that not a single microbe (among *S* species in our pool) that is missing from the current steady state can successfully grow in it, thereby invading the ecosystem and modifying the steady state. To be able to grow, both λ^(*c*)^ and λ^(*n*)^ of the invading microbe has to be larger than those constants for each of the resident microbes (if it exists) currently limited by these resources.

**FIG. 1.**
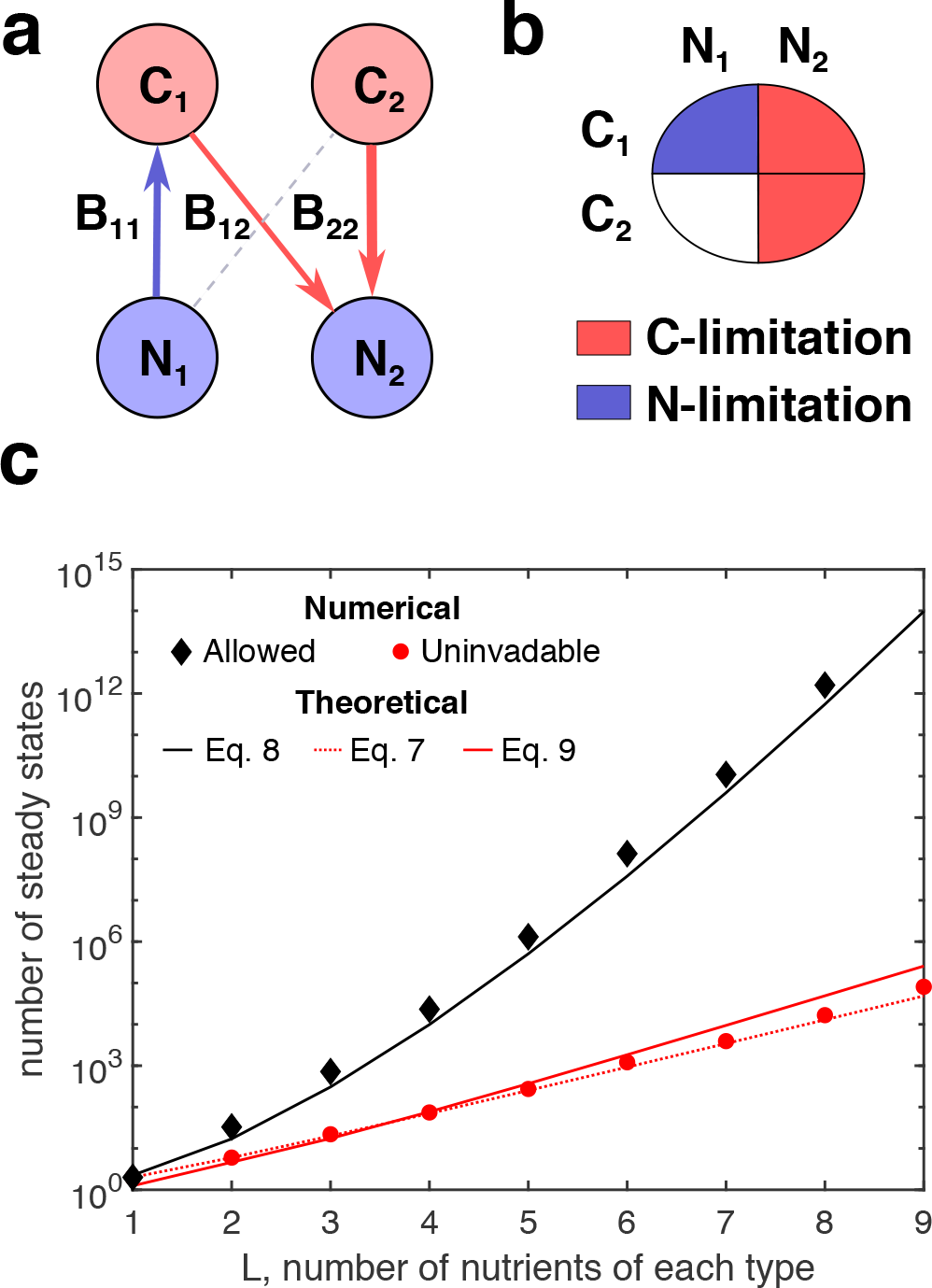
Number of allowed and uninvadable states. (**a-b**) Two equivalent schematic representations of a simple community guided by exclusion rules in our model. (**a**) Bipartite network representation of an allowed state. Nodes of two types represent different nutrients, arrows represent different bacteria species, direction and color of each arrow corresponds to the nutrient limitation of each species (arrow goes from the limiting nutrient to non-limiting one), (**b**) Matrix representation of an allowed state. The allowed state shown in panel A can be represented as a matrix with *K* = 2 rows (representing the 2 carbon nutrients) and *M* = 2 columns (representing the 2 nitrogen nutrients). Cell (*i*, *j*) of the matrix represents the species consuming *i*^th^ carbon source and *j*^th^ nitrogen source. A colored cell implies the presence of the species and an empty cell signifies its absence. If the species is present, it can be limited by either the carbon it is utilizing (in which case the cell is colored red) or the nitrogen it is consuming (colored blue). (**c**) The number of allowed states including both invadable and uninvadable steady states, red squares, and the subset of uninvadable states, black circles, obtained by exhaustive testing of all possible states against the exclusion rules 1 and 2. The x-axis is the number of nutrients of each type (*L* carbon sources and *L* nitrogen sources). The pool has *L*^2^ species - one for each pair of nutrients. Red and black lines are the theoretical estimates for each of the numbers for continuous approximation. Red dashed line is the lower bound to the number of uninvadable states based on the stable marriage model and given by the Eq. 6. Note the logarithmic scale on the y-axis.

### Total number of allowed and uninvadable states

A natural question to ask is how many allowed states and, separately, how many uninvadable states are in principle possible for a given pool of species (each state will be realized for different environmental parameters). Our exclusion rules allow one to identify all of them *based only on the set of ranked tables of competitive abilities of all microbes for each of the nutrients*. For large *K*, *M*, and *S* this is computationally expensive. Indeed, in a brute force method one has to check all of the 3^*S*^ candidate states (each of *S* species could be either absent, or, if present - limited by either carbon or nitrogen) for compliance with the exclusion rules 1 and 2. Each of the allowed states then need to be checked for invasion against up to *S* − 1 missing species to verify their uninvadability.

To help the process of search for allowed and uninvadable states we mapped the problem of finding them to that of finding all stable matchings in the celebrated stable marriage and college admissions problem in game theory and economics^52^, which is algorithmically well-studied^53^. This connection is described in detail in Supplementary Note 3. In a nutshell, we found a one-to-one correspondence between the set of uninvadable steady states in our model and the set of stable matchings in Gale-Shapley college admission problem. One first allocates the number of “partners” (in-degrees in our network representation of steady states) for every resource of a given nutrient type (say N). As described in Methods Section, one can then use the mathematical machinery of the stable marriage problem to discover all stable matchings in which all C sources have out-degree 1 and each of N sources has in-degree prescribed by our selected allocation.

Since the number of ways of selecting the number of partners is exponentially large (the combinatoric factor is shown below), and that for each such distribution the Gale-Shapley theorem guarantees at least one stable matching, the overall number of uninvadable states is also exponentially large and is bounded from below by:

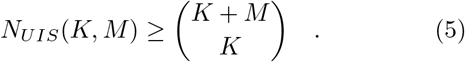

For equal (and large) number of carbon and nitrogen sources *K* = *M* = *L* this estimate can be further simplified to give:

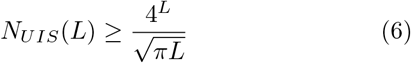

(see Supplementary Note 3 for details). Note that the connection between uninvadable states in our system and the stable marriage problem is rather different than that in Ref.^19^. Indeed, while in Ref.^19^ “stable marriages” are established between microbes and sequentially-used (diauxic) substitutable resources, in the present study the “marriages” are between different sources of carbon and nitrogen, while microbes play the role of “matchmakers” (connectors).

To verify these mathematical results we carried out numerical simulations of the model with equal number - *L* - of carbon and nitrogen sources and a pool of *L*^2^ species, with exactly one microbe using each pair of nutrients. The number of states revealed by our numerical simulations is indeed very large. For example, for only 9 carbon resources, 9 nitrogen resources and a pool of 81 species, the microbial community is capable of 81,004 distinct uninvadable states and roughly 10^14^ allowed states. Fig.1 shows the numerical results, which are in agreement with our theoretical predictions. The number of allowed states increases faster than exponential. In the continuous approximation (solid black curve in Fig. 1C) it is asymptotically described by

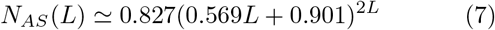

(see Supplementary Note 4 for details). The number of uninvadable steady states also rapidly increases with *L*. While for *L* ≤ 9 it rather closely follows the lower bound given by Eq. 6 (dashed line in Fig. 1C), for larger values of *L* we saw a crossover to a faster-than-exponential regime where it grows as

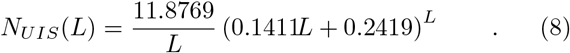

This asymptotic formula derived in Supplementary Note 4 is accurate for values of *L* much larger than those shown in Fig. 1C. However, the numerical integration of the expression derived in the continuous approximation (solid red line in Fig. 1C) is close to the exact number of uninvadable states (see Supplementary Note 4). The discrepancy is likely due to the fact that the continuous approximation assumes that the distribution of the number of species per each pair of resources is Poisson with mean equal to 1 (instead of exactly one species in our numerical simulations). Note that this growth is much faster than sub-exponential expression recently calculated for Lotka-Volterra model with strong interactions^58^. However, unlike Ref.^58^, we calculate the total number of possible stable uninvadable states feasible for different values of environmental parameters.

### Feasible regions of nutrient influxes for each of the steady states

Steady states allowed by Eqs. 2, 3 (satisfying the constraints imposed by our two exclusion rules) are further constrained by Eqs. 4. That is to say, for a given set of the *K* + *M* nutrient influxes only a very small subset of exponentially large number of allowed states would be feasible. From another angle, according to Eqs. 4 each allowed state has a finite region of nutrient influxes where it is feasible. Similar to Ref.^59^ (for Generalized Lotka-Volterra model) and Ref. ^46^ (for consumer-resource MacArthur model) testing if a given Allowed State is feasible at a specific nutrient influx requires inverting the matrix in Eqs. 4 (see Eqs. 9 and 10 in Methods Section) to get the unique set of bacterial populations *B*_*α*_ for all species present in a given allowed state.

In principle one can also find the set of all nutrient influxes for which a given allowed state is feasible in the opposite way that does not involve matrix inversion. One just needs to span (or sample by a Monte-Carlo simulation) the *K* + *M*-dimensional space of positive microbial populations and non-limiting nutrient concentrations in this allowed state. For each point in this set Eqs. 4 trivially define all nutrient influxes required to realize these populations/concentrations in the steady state defined by a given allowed state (see Methods Section for details).

The volume of the region of feasible influxes quantifies the structural stability^60^ of the steady state. States with larger volumes are expected to be more robust in case of fluctuating nutrient influxes. In order for this volume to remain finite, we impose an upper bound on the influx of each nutrient: 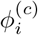, 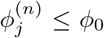. Another way to quantify the structural stability of each allowed state is to calculate the volume of all influxes as bacterial abundances and non-limiting nutrient concentrations vary within a given positive range. Structural stability defined this way is proportional to det 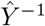, where 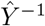 is the matrix of inverse yields by which the vector of bacterial populations is multiplied in the Eq. 4.

Each region of feasible influxes is generally bounded by multiple hyperplanes in a *K* + *M*-dimensional space and thus is difficult to visualize. In Fig. 2A we show feasible volumes of different allowed states in a model with *K* = *M* = 2 nutrients (i.e., two carbon and two nitrogen sources) and *S* = 4 species with exactly one species for each pair of these nutrients. Hereafter we denote this example as the 2C×2N×4S and use similar nomenclature for other examples. For a particular choice of λ-values used in our numerical simulations of the 2C×2N×4S system (see Supplementary Tables I, II for values of λ and yields) we get 33 allowed states (plus 1 empty state without microbes) that do not violate the two rules of competitive exclusion. It is well below 3^*S*^ = 3^4^ = 81 candidate states possible before competitive exclusion rules were imposed. We labeled these allowed states in such a way that the first seven of them (S1-S7) are also uninvadable. Fig. 2A visualizes the feasible region of each state as an ellipse, with its center positioned at the center of mass of feasible influxes, and its area selected to cover 25% of feasible influxes. To better separate the feasibility regions of these states we performed principle component analysis using center of mass for each state. The website given in Supplementary Information attempts a more realistic visualization of the 6 uninvadable states, which are also dynamically stable (see section on dynamical stability below) as linearly confined regions in the 3-dimensional (out of the total of 4 dimensions) PCA space. As one can see from this figure, structural stabilities (feasible volumes) of different states vary over a broad range with S1, S2, and S3 being the most structurally stable with large feasible volumes, while S4, S5, and S6 are considerably smaller (they are the narrow stripes, sandwiched between the three largest states).

**FIG. 2.**
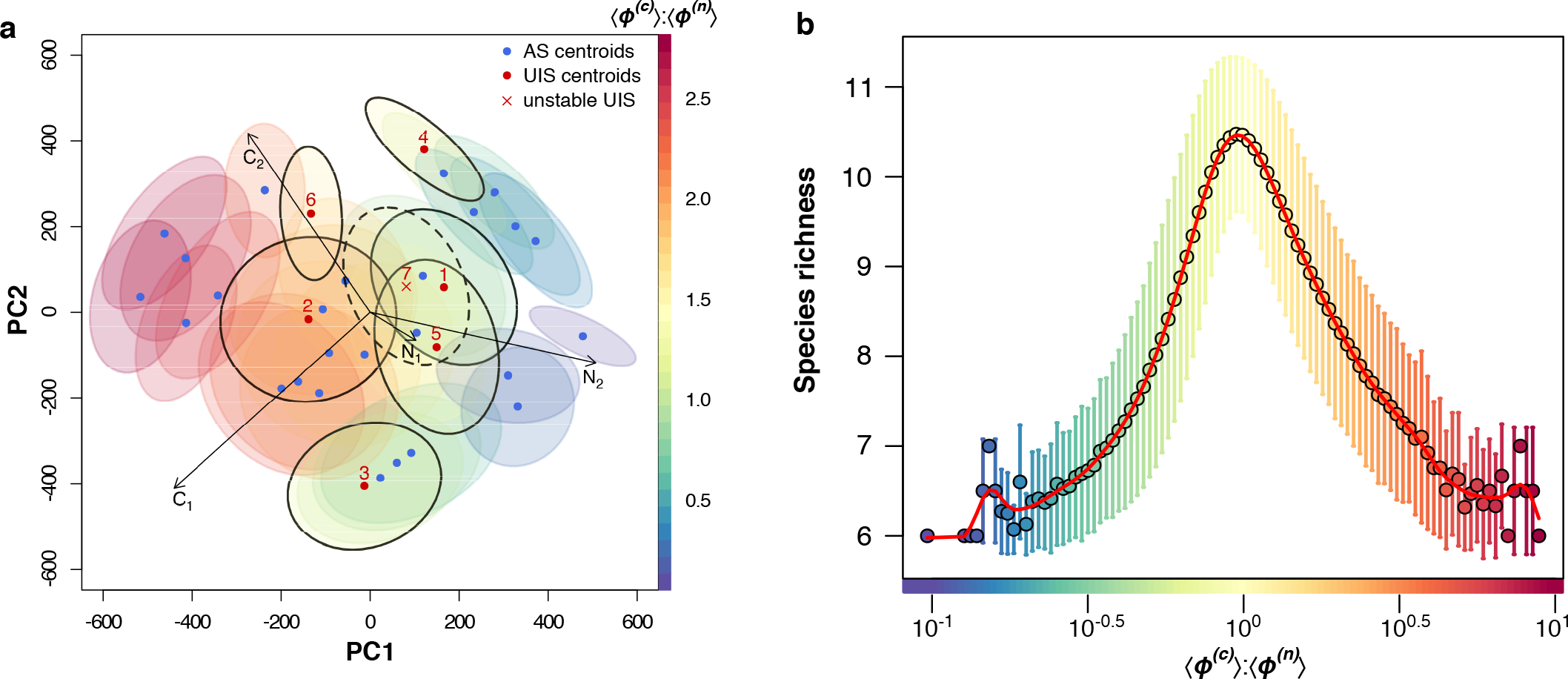
Feasible volume of states and average species richness. (**a**) Principle Component Analysis (PCA) of the 4-dimensional vectors of average nutrient fluxes feasible for the 33 allowed states in our 2C×2N×4S example (see λ and yields in Supplementary Tables I, II). The ellipse around each dot approximates the boundary encompassing 25% of feasible nutrient influxes for each of the 33 allowed state. The arrows in the middle correspond to the direction of changes of 4 nutrient fluxes in PCA coordinates. (**b**) Dependence of *species richness* on the averaged *ø*^(*c*)^: *ø*^(*n*)^ flux ratios for our 6C×6N×36S example (See Supplementary Tables III, IV, V, VI for the values of λ and yields used for this example). Each point represents the average number of surviving species in all the stable uninvadable states which are feasible in an interval of nutrient influxes with average *ø*^(*c*)^: *ø*^(*n*)^ ratio, partitioned into 100 bins. Error bars show standard deviation of the species richness around each interval and the solid red curve is a trend line. Colors on both plots corresponds to similar *ø*^(*c*)^: *ø*^(*n*)^ ratio.

We further explored a more complex model with *K* = *M* = 6 nutrients and 36 species, i.e., 6C×6N×36S (see Supplementary Tables III, IV, V, VI for the values of λ and yields). For this choice of λ we obtained a total of 134,129,346 allowed states out of which 1211 were unin-vadable. Using Monte-Carlo simulations over a region of nutrient influx space (see Methods Section for details) we obtained the feasible volumes of each of the uninvadable states. Utilizing this data we explored how environmental parameters (in our case nutrient influxes) affect the number of surviving species. In Fig. 2B we show that species richness peaks when the fluxes of the two available nutrients are most balanced and it falls with increasing disproportionate between the two.

### Dynamical stability of steady states

So far we avoided an important question of dynamical stability of steady states in our model. We tested the stability of all allowed states in our 2C×2N×4S example by performing computer simulations in which each of the 33 allowed states was subjected to small perturbations of all microbial populations present in a given state (see Methods Section for details). Naturally, an invadable state will be dynamically unstable against introducing small populations of successful invaders, which does not count as its dynamical instability. In our example only one of the states (S7) was found to be dynamically unstable. Interestingly, it was unstable for all combinations of nutrient influxes we tested, while the remaining 32 allowed states were always dynamically stable. This property of our model is different from, e.g. MacArthur model, where stability of a state generally depends on nutrient influxes and concentrations in the environment^44^. For another variant of the MacArthur model all steady states were found to be stable^46^.

For our 6C×6N×36S example the number of allowed states is too large, hence we wish to classify only the 1211 uninvadable states based upon its dynamical stability. Further, to carry out this classification we use a computationally inexpensive algorithm (as against the direct test of stability used for our 2C×2N×4S example) as described below. Since for each set of influxes there must be *at least one dynamically stable uninvadable state* providing the endpoint of model’s dynamics, an unstable steady state can never be alone in the influx space: wherever feasible, it is bound to decay into one of the stable uninvadable states feasible for these environmental parameters. That provides an intriguing way to use influx maps to infer stabilities of states. Indeed, if for a given state one could find a flux region in which no other uninvadable states are feasible - then this state has to be automatically dynamically stable. In our numerical simulations we found that at each specific influx point *V* stable uninvadable states are always accompanied by *V* − 1 dynamically unstable ones. Application of this indirect algorithm on our 6C×6N×36S example revealed 137 dynamically unstable and 1058 dynamically stable states out of the 1211 uninvadable states. This method could not infer the stability of 16 uninvadable states with very small feasible volumes. Using Monte-Carlo simulations we computed the feasible volume of each uninvadable state for our 6C×6N×36S example. We find that the distribution of volumes of all these states is log-normal (for the distribution of log of volume parameters are: *μ* = −8.87 ± 0.06, σ = 2.08 ± 0.04). We also find that the there is no significant difference between distributions of volumes for the stable and unstable uninvadable states (two-sample Kolmogorov-Smirnov test: *p* – *value* = 0.94).

**FIG. 3.**
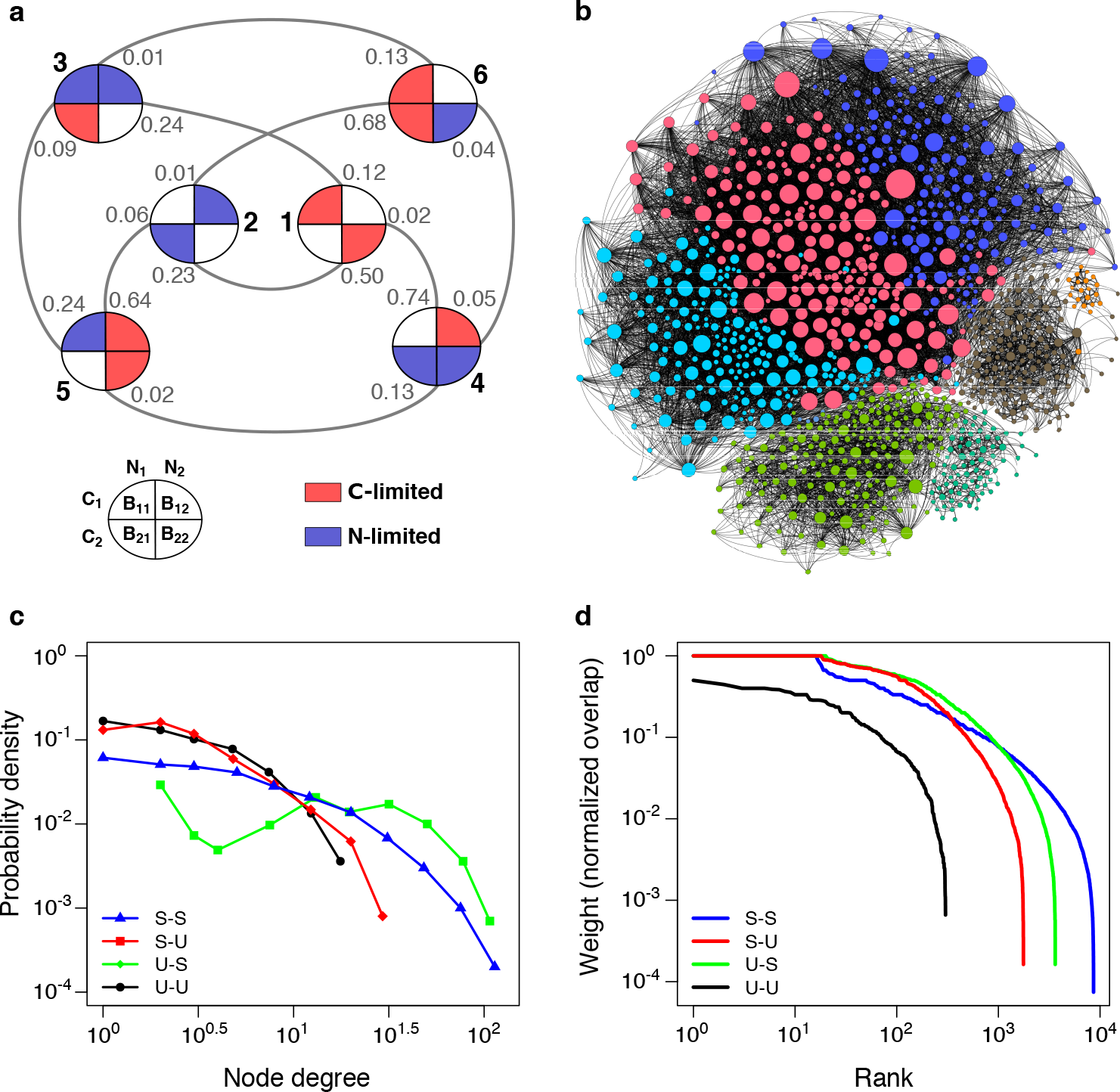
Networks of overlaps between feasible volumes of uninvadable states. (**a**) The network of overlaps among 6 stable uninvadable states in the 2C×2N×4S example (same as in Fig. 2A). Each state is represented in its matrix configuration (see Fig. 2A). A link between 2 states represents an overlap of their feasible volumes. The fraction of the volume of each state that overlaps with another state is the weight of a directed edge indicated on the link connecting these states. Hence, the sum of all these fractions denoted on the links near a state indicates the total fraction of its volume that overlaps with all the other uninvadable state. (**b**) The network of overlaps of the 1195 uninvadable states (both stable and unstable) in our 6C×6N×36S example (same as in Fig. 1B). Nodes and links are the same as in panel a). Size of a node reflects its degree (i.e., the total number of states it overlaps with). The color of each node corresponds to the modularity cluster/class (8 in total) it belongs to (see Methods Section for details). (**c**) Degree distribution of the network in panel (b) with different colors representing different types of degrees: the number of stable states neighboring each of the stable nodes (blue triangles), the number of unstable states neighboring each of the stable nodes (red squares), the number of stable states neighboring each of the unstable nodes (green diamonds), the number of unstable states neighboring each of the unstable nodes (black circles) (**d**) Rank ordered distribution of weights of the network from panel (b). The weights normalized as in (a) represent normalized overlaps of states. Different lines represent overlaps between pairs of two different types of nodes - stable (S) and unstable (U) with the same colors/labels as in panel (c).

### Multistability of microbial ecosystem, alternative stable states for the same environmental parameters

Our model is generally capable of bistability or even multistability when two or more uninvadable states are feasible for the same environmental conditions given by nutrient influxes. This happens when the feasibility regions of multiple stable uninvadable states overlap with each other. For influxes in the intersection area, all of the overlapping states are feasible and, since each of them is uninvadable, they cannot transition to each other through addition of other species from the pool. Fig. 3A shows the network of such overlaps between the 6 stable uninvadable states in our 2C×2N×4S example. The fractional number shown on each edge represents the fraction of the feasible volume of each state, over which it overlaps with its neighboring state.

The network of overlaps of the 1030 uninvadable states (both stable and unstable) that have at least one neighbor in our 6C×6N×36S example (same as in Fig. 2B). Nodes and links are the same as in panel a). Size of a node reflects its degree (i.e., the total number of states it overlaps with).

In Fig. 3B we plot the network of overlaps between feasible volumes of the 1195 uninvadable states in our 6C×6N×36S example. We performed the standard modularity analysis on this network (see Methods Section for details) and obtained 8 clusters indicating that the states are not randomly distributed but clustered in the flux space. We distinguished two types of nodes in this network - dynamically stable (S) and unstable (U) ones. Thus in Fig. 3C we distinguish between four different types of degrees: the number of stable states neighboring each of the stable nodes (blue triangles), the number of unstable states neighboring each of the stable nodes (red squares), the number of stable states neighboring each of the unstable nodes (green diamonds), the number of unstable states neighboring each of the unstable nodes (black circles). One can see that all four type of degrees vary over a broad range with the largest degrees of the four types listed above equal to 164, 41, 115, and 21 correspondingly. Thus the biggest hub among stable states is connecting to around 15% of other stable states. Rank-ordered distribution (Zipf plot) of edge weights of the network from Fig. 3B are are shown in Fig 3D. Different colors and symbols represent overlaps between different types of nodes (S-S, S-U, U-S, and U-U). Here the x-axis shows the rank of the weight of a certain type (1 being the largest) and the y-axis shows the value of this weight. Figure 3D shows that different types of edges have different probability distributions of weights (all of them broad).

We can define *coexisting* states as a set of states which are *simultaneously* feasible in a finite region in the nutrient influx space. This determines the multistability of the system. The number of such coexisting stable uninvadable states never goes above 2 in our 2C×2N×4S example with λ and yields as in Fig. 3A. However, for a larger number of resources we do observe multistability of more than 2 stable uninvadable states. E.g., for our 6C×6N×36S example (with λ and yields same as chosen for Fig 3B) we notice up to 5 coexisting stable uninvadable states (see bold black line in Fig 4B). It is still a far cry from an exponentially large number of all uninvadable states possible for different combinations of nutrient influxes. Also notice that the volume of the influx space occupied by *V*-stable states falls of with *V* faster than exponentially.

We further explored the factors that determine this multistability. Like in its simpler special limit studied in Ref.^47^, the multistability in our model is only possible if individual microbial species have different C:N stoichiometry, quantified by the ratio of their carbon and nitrogen yields. Our numerical simulations (see Methods Section for details) strongly support that when all species have the same stoichiometry 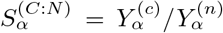, for every set of nutrient fluxes there is a unique uninvadable state. The same is true in the MacArthur model provided that the ratio of yields of different nutrients is the same for all microbes^61^. Simulations on the 2C×2N×4S example (see Fig. 4A) supports our claim that the more narrow is the spread of 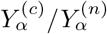 oin our pool of species, the smaller is the average volume of multistable states realized among all possible combinations of nutrient fluxes. Interestingly, in our experiments with varying stoichiometry roughly a half of yield combinations resulted in multistability (i.e., out of 4000 numerical experiments we performed, only for 2069 a non-zero fraction of multistability was observed).

We also simulated our 6C×6N×36S system (with the same values of λ) for many different sets of yields. Fig 3B shows the fraction of nutrient influx space that permits the coexistence of *V*-stable states for 3 different sets of yields, one of which (as mentioned above) permits multistability up to 5. The other two sets of yields show a smaller multistability of 3 and 4 uninvadable stable states.

### Colonization dynamics

In the course of sequential colonization by species selected from our pool (see Methods Section for details), microbial ecosystem goes through a series of transitions between several feasible allowed states culminating in one of the uninvadable states. The set of all colonization trajectories can be visualized as a directed graph with edges representing transitions caused by addition of species from the pool to each of the state. Both the set of nodes (selected among all allowed states) as well as the set of possible transitions between these nodes are determined by the environmental variables (nutrient influxes).

In Fig. 5 we show two examples of state transition graphs in our 2C×2N×4S case. For one set of environmental parameters shown in Fig. 5A, our model has 6 feasible and dynamically stable allowed states (plus 1 empty state) connected by 12 transitions triggered by species addition. For the same species pool, changing nutrient fluxes (Fig. 5B) results in a different set of 10 feasible allowed states connected by 19 transitions. The uninvadable states are visible as terminal ends of directed paths in the top layer of Figs. 5A, B. While the fluxes used in Fig. 5B allow for only one uninvadable state, S5, a different set of fluxes used in Fig. 5A permit for two alternative uninvadable states, S1 and S2. Which of these two states is realized depends on the order in which species were added to the system.

**FIG. 4.**
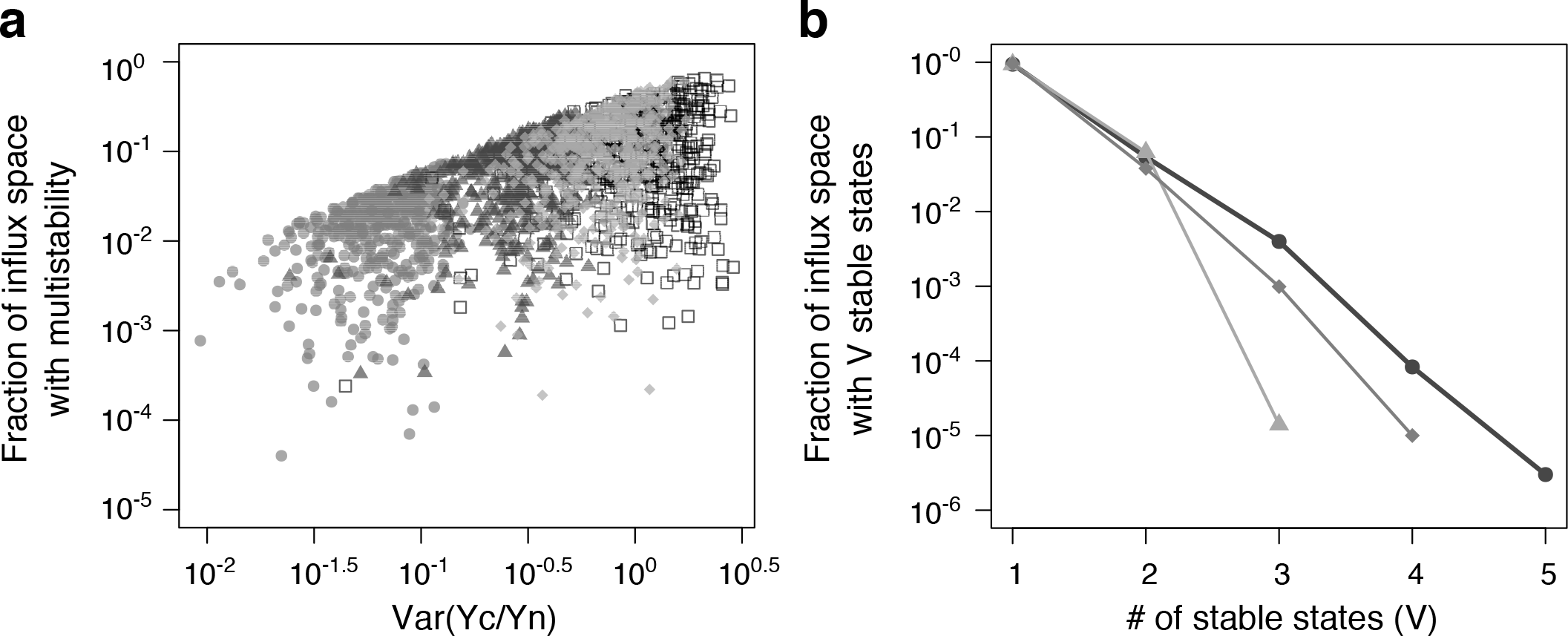
Statistics of multistability for different combinations of yields. (**a**) Fraction of the nutrient influx space with multistability for different set of yields in the 2C×2N×4S example (λ are the same as in Fig. 2A). Y-coordinate of each point represents the fraction of the nutrient influx space (out of 10^5^ influx points we sampled) where multistability was observed for a given set of yields (while keeping competitive abilities of all species fixed). Its x-coordinate represents the variance between the ratios of yields (*Y*^(*c*)^/*Y*^(*n*)^ across all species. In total, 4000 numerical simulations were done, each with a different set of yields. The 4 different types of points indicate the 4 different (uniform) distributions (with different variance) from which the yields of different species were chosen (see Methods Section for details). (**b**) For 3 different sets of yields we plot the fraction of nutrient influx space with V stable uninvadable states for the 6C×6N×36S model (λ are the same as in Fig. 2B). For each set of yields we explored the fraction of the nutrient influx space (out of 10^6^ influx points) that contains V=1,2,3,4,5,.. stable uninvadable states. The figure shows that a difference in the yields of the species results in difference in multistability, although the values of λ (and hence the set of uninvadable states) are the same in all the three cases. Bold black curve corresponds to the example shown in Fig. 3B.

As shown in Fig. 5C, D transitions between these and other alternative states can also be triggered by changing nutrient fluxes. Whenever multistability is present, transitions happen in a hysteretic manner. In Fig. 5C we show a full cycle of changing fluxes, first up (grey line) and then down (black line) in, which results in a system going through a series of discontinuous transitions between states and ending up in a different uninvadable state than that it started from. Where as in contrast, in Fig. 5D there is a unique uninvadable stable state feasible at a certain flux point. This matches several hallmark properties of alternative stable states defined in Ref.^18^ as having “discontinuity in the response to an environmental driving parameter”, lack of recovery after a perturbation (hysteresis), and “divergence due to different initial conditions”.

## DISCUSSION

Ever since Robert May’s provocative question “Will a large complex system be stable?”^23^ the focus of many theoretical ecology studies has been on dynamical stability of steady states in large ecosystems. Unlike the classic MacArthur model^46^, but similar to the original Tilman model^47^, our model is characterized by a mixture of dynamically stable and unstable states. Based on a small sample of examples that we analyzed in detail, we found that the stable states in our model generally outnumber the unstable ones. For example, in the 2C×2N×4S example used above only one state out of 33 is dynamically unstable, while for the 6C×6N×36S example used in Fig 2C and Fig 4, we found only 137 unstable states out of 1195 uninvadable states that we were able to classify using our methods. Another interesting observation is that for randomly selected carbon and nitrogen yields of individual species (defining its C:N stoichiometry) with probability around 50% one ends up with an ecosystem lacking both unstable states as well as alternative stable states.

In fact, the existence of dynamically unstable states in our model always goes hand in hand with multistability. Indeed, in the simplest case of a bistable ecosystem considered in Ref.^15^, a single dynamically unstable steady state always separates two stable states of an ecosystem. Depending on perturbation this unstable state would collapse to either one of the two alternative stable states realized for the same environmental parameters. Interestingly, in our model we always found *V* − 1 unstable states coexisting with V alternative stable states for the same environmental parameters. While, this result is natural for dynamics maximizing a *one-dimensional* Lyapunov function where *V* maxima (corresponding stable states) are always separated by *V* − 1 minima (unstable stable states), we currently do not understand why this rule seems to apply to our *multi-dimensional* system. Indeed, in 1D any smooth function bounded from above and reaching −∞ at *x* = ±∞ always has *V* maxima and *V* − 1 minima. In higher dimensions this property imposes additional constraints on indices on critical points other than maxima which are dictated by the Morse theory^62^. We leave the search for this Lyapunov function for future studies.

**FIG. 5.**
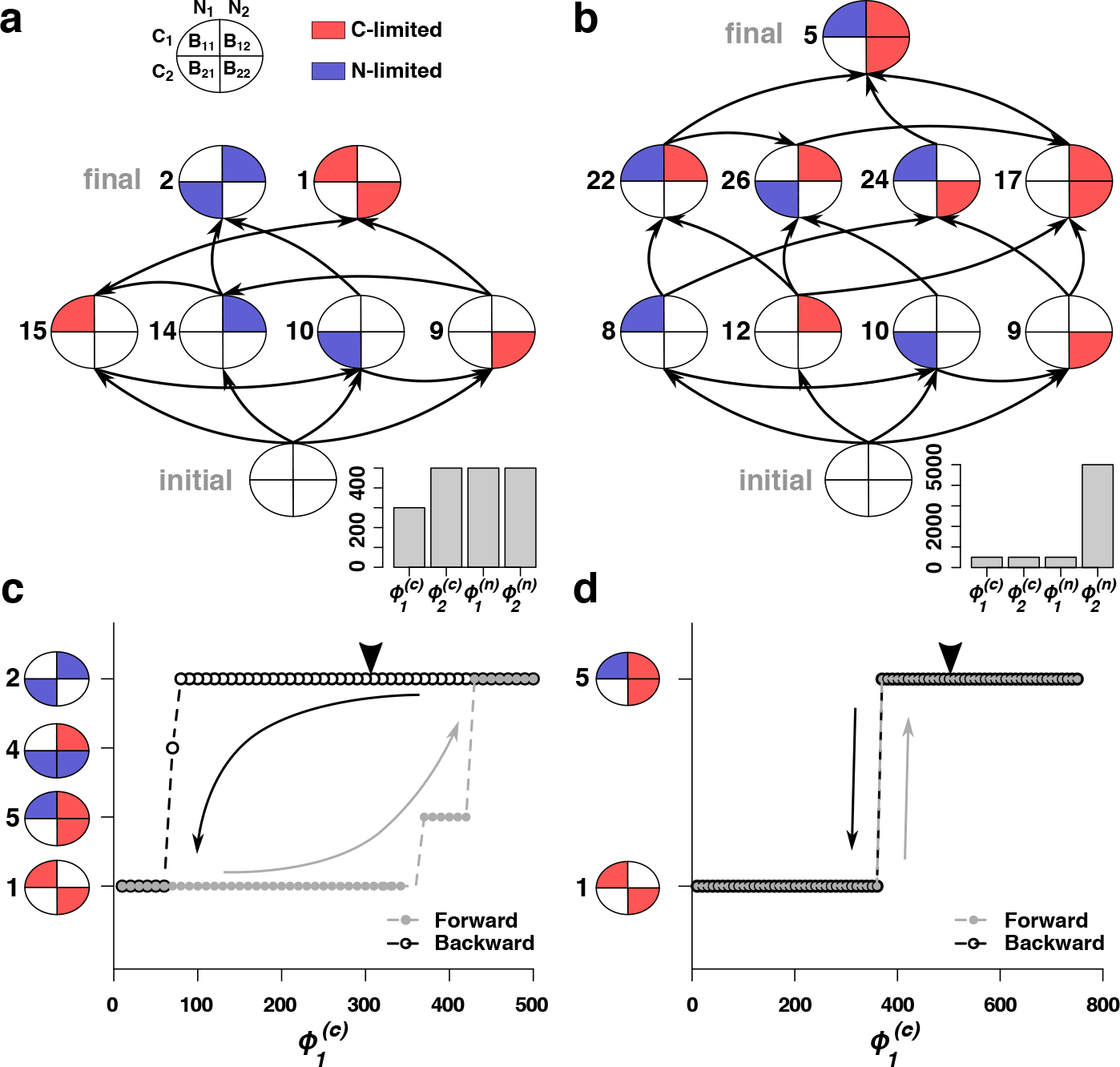
State transitions triggered by colonization dynamics and changing environment in 2C×2N×4S example. (**a**, **b**): Transitions triggered by colonization dynamics. We show the graph of all possible state transitions for our 2C×2N×4S example (same as in Fig. 2A) for two different sets of environmental parameters shown as barplot in **a** and **b** (see Methods Section for details). Starting from an empty set (i.e., no bacterial species) we randomly select and introduce species from the pool (one at a time) and wait for the system to settle into a dynamical steady state, and this process is repeated until the system reaches an uninvadable state. (**c**, **d**): Transitions triggered by changing environment. Here we show the transition between states when one of the parameters 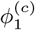 is modified from its original value (pointed black arrow) in panel **a** and panel **b** (shown in panel **c** and panel **d** respectively). (**c**): The environment is set to a low 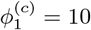 units, with other influxes same as in panel **a**) and on executing colonization dynamics the system settles to the uninvadable state S1. We next increment 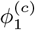 by a unit of 10 and apply colonization dynamics on the current state of the system and iterate this process until 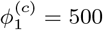. Grey filled dots indicate the states that the system experiences at the end of each colonization dynamics on its journey from 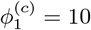 to 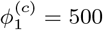. One can see that at 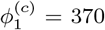 the system transitions to the uninvadable state S5 and at 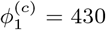 it jumps to state S2 and stays there up till 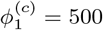. Starting from this state (S2) the above process was repeated but with decreasing 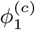. Black empty circles highlight the states observed by the system until 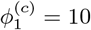. This completes a full cycle of changing 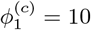. It can be seen that in the transition from state S2 to state S1 occurs at a much lower value of 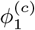 compared to the transition from S1 to S2 when 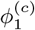 was increased, hence displaying the phenomenon of hysteresis. (**d**): Similarly to panel **c** we varied 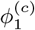 from a low to high value, keeping all the other nutrient influxes same as in panel **b**. In contrast to **c**, we do not observe any hysteretic transitions when 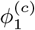 is changed in the forward and reverse directions. The black arrow points at the environmental conditions from panel **b**.

As we demonstrated above, both unstable states and multistability are possible only if different microbes have different C:N stoichiometric ratios. The same is true for models in which microbes co-utilize multiple substitutable (say carbon) nutrients. Indeed, a convex Lyapunov function defined in Ref.^41, 63, 64^ guarantees that for any set of nutrient influxes there exists exactly one stable equilibrium. While the standard MacArthur model has no multistability and, as proven in Ref.^46^, all of its steady states are dynamically stable, its variant in which different microbes have different yields on the same nutrient, has both these properties^61^. Different yields of different microbial species prevent one from constructing the Lyapunov function used in the standard MacArthur model^41^. We leave the topic of existence and the functional form of the Lyapunov function in our model for future studies.

Using the algorithms of the stable matching problem^52, 53^ we were able to list all of these states based only on ranked tables of nutrient competitive abilities of different microbes. The advantage of this approach is that it bypasses the need for precise measurements of the kinetic parameters and depends only on the relative microbial preferences toward nutrients. This property is likely unique to our model. Indeed, for a popular MacArthur model^40, 41, 64^ of co-utilization of fully substitutable resources, the relative rankings of different microbes for nutrients depend on nutrient concentrations. This greatly complicates the task of deducing the ultimate set of allowed states, that is to say, all subsets of surviving microbes realized for different environmental conditions.

To improve mathematical tractability of our model we have made a number of simplifying assumptions. These can be relaxed in the following variants of our basic model some of which are listed below: (i) A simple generalization of our model is to relax the condition of extreme specialization to allow for generalist species, i.e., those with growth rate is given by:

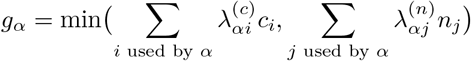

Here the sum over *i* (respectively *j*) is carried out over all carbon (respectively nitrogen) sources a given species is capable of using. Here one assumes that multiple sub-stitutable sources are co-utilized as in the MacArthur model. Another possibility is to assume that each species is using its substitutable resources one-at-a-time, as we assumed for multiple carbon sources in Ref.^19^. Since at any point in time each of the species is using a “specialist strategy” growing on a single carbon and a single carbon source, we expect many of our results to be extendable to this model variant. Using either of these model variants one can explore environmental conditions (number of resources and their fluctuations in time and space) that would favor specialists or generalists over each other. It will also be interesting to explore how the presence of generalists affects the number of alternative stable states and how the available nutrients are partitioned between the different coexisting species in such a community. (ii) We worked with a fixed size of species pool for our models. By relaxing this constraint and having a large universe of species to choose from, one can explore interesting aspects of evolution and adaptations in our model setup, e.g., how the presence of multistability affects the process of community assembly. (iii) One can also introduce cross-feeding between the species, thus generating additional resources in the system and allowing for a larger number of species to coexist, and, hence further increasing the number of alternative stable states. (iv) Although we have identified that larger variance in C:N stoichiometry of individual species (quantified by 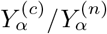 in our model) promotes multistability, other factors affecting the likelihood of alternative stable states remain to be identified in future studies.

## ACKNOWLEDGMENTS

Part of this work has been carried out at the University of Padova, Italy, in August 2018, during a scientific visit by one of us (S.M.).

## AUTHORS CONTRIBUTIONS

S.M. designed the research; P.P. simulated the computational model; Y.F. and S.M. developed the theory for the computational model; P.P., S.M., and V.D. analyzed the data; S. M. V. D. and P.P wrote the manuscript; and S.M. supervised Study.

## METHODS

### Enumeration of all allowed states

Every allowed state can be converted into a unique bipartite network with the two types nutrients (carbon and nitrogen) as the nodes and links representing the surviving species. The source node of the link represents the rate limiting nutrient of the species and each link is characterized by its λ^(*c*)^ and λ^(*n*)^. (see Fig. 1A). a, The competitive exclusion rules int the language of networks is stated below:

■ **Rule 1** - All nodes can have at most one outgoing link
■ **Rule 2** - All incoming links at any node should have a larger λ than the λ of the outgoing link (if any).

To obtain all allowed states one needs to perform an exhaustive search of all the networks that satisfy the above constraints. We do this in the following way: We start with choosing a particular set of ranked values of 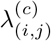 and 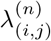 for all *L*^2^ species in the pool. We then choose any one type of nodes on which the outgoing links will be assigned first (say carbon; the choice carbon as opposed to nitrogen does not affect the final result of this algorithm). We then perform a two-step procedure for links allocation. We first allocate all outgoing links from C-type to N-type nodes by choosing a rank between 1 and *L* + 1 for each of the C-type nodes, where rank *L* + 1 corresponds to having no link (no species is limited by this nutrient). For this specific combination of outgoing links we generate all allowed sets of incoming links (from N-type to C-type). To implement this step one needs to follow *Rule 2* to filter out prohibited allocations of incoming links. This procedure is guaranteed to find all allowed states for this chosen specific chosen set of outgoing links from C and one of the allowed states will be uninvadable (see Supplementary Note 3 for details). One repeats this allocation procedure for each combanation ranks of outgoing links for the C-type nodes ((*L* + 1))^*L*^ possible allocation in total) to get a list of all allowed bipartite networks for the set of 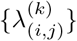

We used the above procedure to enumerate allowed states for different numers of resources *L* (see Fig. 1C). The values of 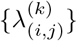 were chosen randomly between 10 and 100 for these numerical experiments.

### Feasibility of the allowed states

As described in the main text, at the steady state of each allowed state the concentration of the surviving species and and the concentration of the nutrients not limited by any of the surviving species is completely determined by the *K* + *M* mass conservation laws (Eqs. 4). Hence each allowed state can be uniquely characterized by a set of *K* + *M* variables (we define them as *X*_*p*_ for the *p*^*th*^ state) consisting of the population of the *S*_*surv*_ ≤ *K* + *M* species and the concentration of the *K* + *M* − *S*_*surv*_ non-limited nutrients. E.g., for the 2C×2N×4S case used in the main text in S5 we have: *X*_5_ = {*B*_(1,1)_, *B*_(1,2)_, *B*_(2,2)_, *n*_2_}.

Now, given the parameters defining the species (i.e., λ and *Y*) and the chemostat dilution constant *δ*, each state *p* will have a finite region in the nutrient influx space (a *K*+*M* dimensional space 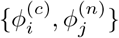)this state will be feasible, i.e., *X*_*p*_ > 0. The volume of this region quantifies the structural stability of the state *p*.

To simplify the process of calculating the feasible volumes we assumed the *high-influx limit*, i.e., 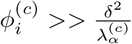 and 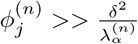. It means that if any nutrient is limited by a surviving species in the allowed state, the concentration of that nutrient at the steady state will be negligible compared to what it was before speciation took place. With this reasonably valid *high-influx limit* assumption the mass conservation laws (Eqs. 4) that are used to obtain the feasible volumes of each of the allowed states can be represented into a compact matrix form. For the allowed state *p* the matrix equation becomes:

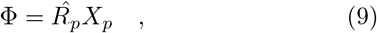

where Φ is the vector of the *K* + *M* nutrient fluxes and 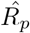 is a matrix composed of *Y*^−1^ of surviving species and “1” for each of the non-limiting nutrients in the allowed state *p*. For the S5 in 2C×2N×4S example the Eq. 9 expands to:

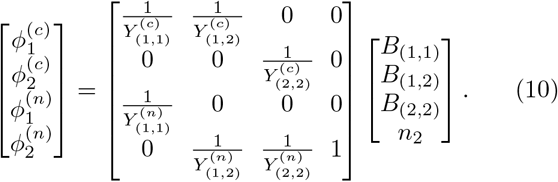

### Monte-Carlo sampling of nutrient influx space

Using Eq. 9 it is trivial to compute if an allowed state is feasible at a particular nutrient influx point Φ. To check feasibility of the allowed state *p* at Φ we simply multiply the inverse of the matrix 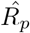 with the vector Φ. If all the elements of the resulting vector *X*_*p*_ are positive then the allowed state *p*is feasible at Φ. If the matrix 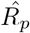 is not invertible i.e., 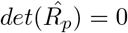, it indicates that this allowed state is not feasible anywhere in the nutrient influx space.

We imposed a common upper and lower bound on each of the *K* + *M* nutrient influxes 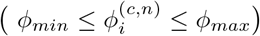 thus restricting the search of volumes of feasible allowed states in a *K* + *M* dimensional hypercube in the nutrient influx space. We chose ø_*min*_ = 10, ø_*max*_ = 1000. The lower bound ensures that the system is always in the *high-flux limit* as 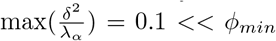. This is because *δ* =1 and λ_**min**_ = 10 (λ were chosen uniformly between 10 and 100). We then randomly spanned a million points in this hypercube and checked feasibility of each allowed state (i.e., for state *p* we checked that all the elements of *X*_*p*_ are positive). For each state we calculated the total number of points where it was feasible to quantify the volumes of the allowed states.

### Overlap of Volumes

Two allowed states are said to overlap with each other if there exists a set points in the hypercube at which both of them are feasible. We used the data obtained by Monte-Carlo sampling to calculate shared feasibility regions (shared sets of points) between states. We then normalized those numbers for each state by its overall volume (see Fig. 3A). For 6C×6N×36S case we used Gephi 0.9.2 software package to visualize the network and performed modularity analysis to identify densely interconnected clusters in the network^65^. Resolution parameter was set to 1.5 to produce Fig. 3B.

### Dynamical stability of the allowed states

We checked the dynamic stability of the allowed states in two ways:

1. **Perturbation analysis**. We prepared each allowed state at one of its feasible influx points and subjected it to small perturbations of (i) all the *K* + *M* = 2 nutrient concentrations and (ii) the populations of the *S*_*surv*_ species in the state. The importance of perturbing the population of only the *S*_*surv*_ species should be noted because an invadable state, by definition, will always be dynamically unstable against addition of (at least one) new species from the species pool. And this instability should not render the invadable state as dynamically unstable. Hence we stress that an allowed state will be dynamically unstable if perturbation of any of the nutrient concentrations or the population of any of the *S*_*surv*_ species drives the state to a different allowed state.
2. **Overlap analysis**. The dynamic stability of the allowed states can also be inferred from the influx map (a map that gives us the information of all the feasible states possible at each point in the influx space) obtained from our Monte-Carlo simulations. We first recognize all states which had a unique presence at at least one influx point (i.e. no other states are feasible at this flux point). All such states should be dynamically stable by definition. Note that Monte-Carlo samples a finite number of influx points and thus it is possible to miss crucial influx points which could have rendered some of the states as stable and thus leading to assigning some of the stable states as unstable. This false assignment will lead into the violation of the V/V-1 rule (as described in the main text) at some influx points. To correct for this error we go over each unstable state and check if assigning it as stable reduces the list of violated influx points. If it does then we include it in our list of stable states. This method could not infer the stability of 16 uninvadable states which had very small feasible volumes.

### Yield-variation

Since the dynamical stability and the size of feasible volume of the states depends on the choice of values of the yields *Y*, we performed a set of Monte-Carlo sampling experiments for 2C×2N×4S and 6C×6N×36S examples to explore how the choice of yields governs multistability. For 2C×2N×4S case we performed 4000 Monte-Carlo simulations for a fixed set of λ (see Supplementary Table 1) and yields were drawn from uniform distributions with different standard deviations (1000 simulation per standard deviation). We used these simulations to calculate the fraction of influx space where we observed multistability (Fig. 4A). For 6C×6N×36S example (see λ in Supplementary Tables 3-4) we chose 4 different sets of Yields and performed Monte-Carlo simulations for each of them. We further performed overlap analysis for each of these numerical experiments counting the number of uninvadable stable states feasible for each flux point (see Fig. 4B).

### Colonization dynamics

To study the process of speciation in our model, we implemented the sequential colonization procedure as described below. We first set the system at the abiotic state (i.e., 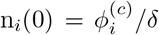 and 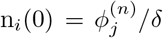) We then randomly select one species from our pool of S species and introduce it into the system with a small population density (10^−5^). We then perform a numerical integration of the current system until the system settles into a steady state. If the population density of any of the species at the steady state falls below a predefined threshold (10^−7^) we considered it to be extinct. We keep performing this random selection and introduction of species addition followed by dynamic integration until no new species from the pool can invade, thus giving us an uninvadable state. This colonization dynamics is repeated for a large number of random-order-introduction of species to obtain all possible terminal ends. The set of all steady states obtained in this process are all the allowed states that the system navigates through.

We followed the above procedure for two different sets of environmental parameters in our 2C×2N×4S example. First set: 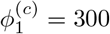, 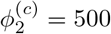, 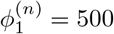, 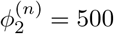, *δ* =1 (see Fig. 5A). Second: 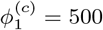, 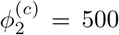, 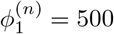, 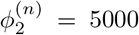, *δ* =1 (see Fig. 5B). This colonization dynamics was repeated for a large number of random order introductions of species from the pool to obtain all possible transitions (shown as black arrows in Fig. 5A, B) between the allowed states at the given nutrient influx.

To study transitions between uninvadable states in response to environmental perturbations, we started from one of the uninvadable states and varied one of the fluxes in some range 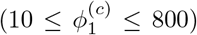 with some step 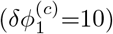 while constantly introducing the random bacterial species from the pool to the system.

The numerical integration for the above process was done in C programming language using the **CVODE** solver library of the **SUNDIALS** package^66^ downloaded from the website: https://computation.llnl.gov/projects/sundials/sundials-software.

## SUPPLEMENTARY INFORMATION

### Supplementary Note 1. General form of growth laws

It is straightforward to generalize our model to include more general functional form for growth laws than Liebig’s law, 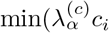, 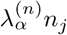. Microbial growth on two essential substrates is thought to normally follow the Monod’s equation for the rate-limiting nutrient: 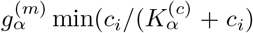, 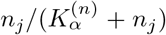 (See Ref.^67^ for a discussion of limitations of Monod’s law). For low concentrations of the rate-limiting nutrient, say carbon source, the Monod’s law simplifies to the proportional growth law used throughout this study: 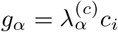. Microbes’ competitive abilities, also known as their specific affinities towards each substrate, are related to the parameters of Monod’s law via

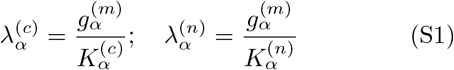

**FIG. S1.**
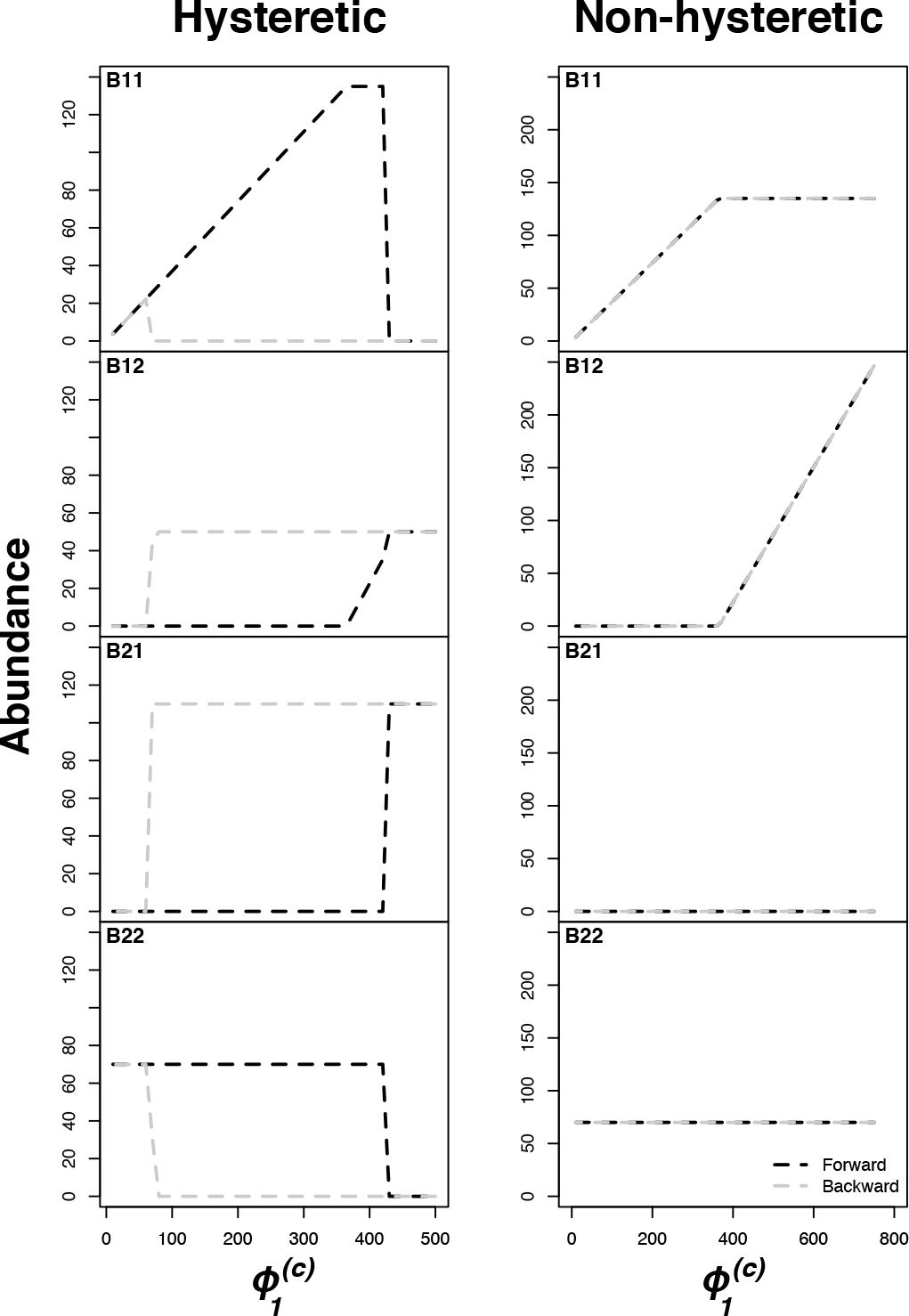
State transitions triggered by colonization dynamics and changing environment in 2C×2N×4S example. Changes in steady state bacterial abundances through flux variation in Fig. 5 (left panel corresponds to hysteretic case in C, right - non-hysteretic transition from D).

In another variant of growth laws, two essential nutrients at low concentrations jointly affect the growth rate of the microbe: 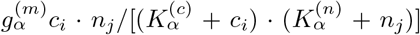 (see Ref.^68^ for a discussion of these and other double-substrate growth laws). For simplicity of mathematical calculation we limit this study to Liebig’s law. However, many of the essential results we obtained (e. g. multistability phenomenon that can be observed for some ecosystems characterized by specific sets of parameters) hold for all the growth laws listed above. In fact, the low concentration version of the previous growth law, where 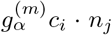 has been studied by one of us in context of autocatalytic growth of heteropolymers^69^. The results of this paper are largely consistent with the present study, namely, in both cases the system has a large number of steady states with between min(*K*, *M*) (corresponding to *Z* in the notation of Ref.^69^) and *K* + *M* − 1 (corresponding to 2*Z* − 1). Regarding why the number of bacterial species can not be larger than the total number of nutrients minus 1, one can prove for any form of the growth laws, that when all yields are the same, states with *K* + *M* species have zero feasible volume, that is to say, they are only possible on a lower-dimensional manifold in the (*K* + *M*)-dimensional space of influxes (this results has been already discussed by Tilman in his special case^47^. Multistability is also possible in the variant of the MacArthur model^40, 41, 64^ in which different species can have different yields when growing on the same nutrients. A convex Lyapunov function^41^ precluding multistability does not exist in this case. We leave this topic for future studies.

### Supplementary Note 2. Constraints on steady states from microbial and nutrient dynamics

A steady state of equations describing the microbial dynamics (Eqs. 2) is realized when either *B*_*α*_ = 0 (the species was absent from the system from the start or subsequently went extinct) or when its growth rate *g*_*α*_ is exactly equal to the chemostat dilution rate *δ*. This imposes constraints on steady state nutrient concentrations with the number of constraints equal to the number of microbial species present with non-zero concentrations. Since, in general, the number of constraints cannot be larger than the number of constrained variables, no more than *K* + *M* of species could be simultaneously present in a steady state of the ecosystem. For Liebig’s growth law used in this study, each resource can have no more than one species for which this resource limits its growth, that is to say, which sets the value of the minimum in 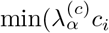, 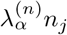 The steady state concentrations of these resources are given by 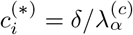 (if the growth is limited by the carbon source) and 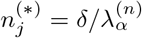 (if the growth is limited by the nitrogen source). Here *α* is the species whose growth is rate-limited by the resource in question. In a general case, no more than one species can be limited by the same resource (carbon in our example), since the species with the largest λ^(*c*)^ would out-compete other species with smaller values of λ^(*c*)^ by making the steady state concentration 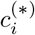 so low that other species can no longer grow on it. Note however, that multiple species *β* could consume the same resource as the rate-limiting species *α*, as long as their growth is not limited by the resource. Each of these species must then be limited by their other nutrient (a nitrogen source in our example). However, their survival requires that carbon concentration set by species α is sufficient for their growth. Thereby, any species growing on a resource in a non-limited fashion must have 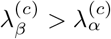.

Mathematically, it cane be proven by observing that, since species *β* is limited by its nitrogen resource, one must have 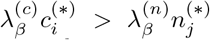. At the same time in a steady state, the concentrations of all rate-limiting resources are determined by the dilution rate *δ* via 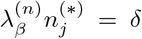, and 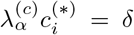. Combining the above three expressions one gets: 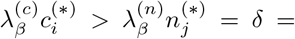 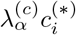, or simply 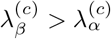. The constraints on competitive abilities A for species present in a steady state in our model are then:

- Exclusion Rule 1: Each nutrient (either carbon or nitrogen source) can limit the growth of no more than one species *α*. From this it follows that the number of species co-existing in any given steady state cannot be larger than *K* + *M*, the total number of nutrients.
- Exclusion Rule 2: Each nutrient (e.g. specific carbon source) can be used by any number of species in a non-rate-limiting fashion (that is to say, where it does not constrain species growth in Liebig’s law). However, any such species *β* has to have 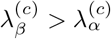, where 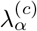 is the competitive ability of the species whose growth is limited by this nutrient. In case of a nitrogen nutrient, the constraint becomes 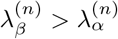.

Note that the steady state solutions of equations 2 do not depend on populations *B*_*α*_ of surviving species. Their steady state populations 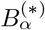 are instead determined by Eqs. 3. Taking into account that, in a steady state, the growth rate of each surviving species is exactly equal to the dilution rate *δ* of the chemostat, after simplifications one gets:

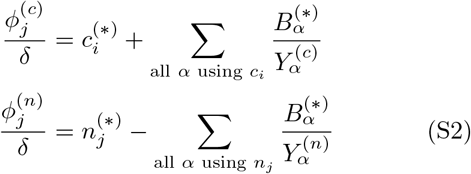

As described above, the steady state concentration of resources are given by 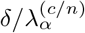, where *α* are the species rate-limited by each resource. In the absence of such species, the concentration of a resource is given by anything left after it being consumed by surviving species in a non-rate-limiting manner. One can show that in this case, the resource (e.g. carbon) concentration has to be larger than 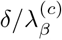, where 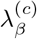 is the smallest affinity among microbes utilizing this resource.

One convenient approximation greatly simplifying working with equations S2 is the “high-flux limit” in which 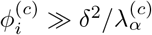 and 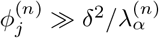. In this approximation one can approximately sets to zero the steady state concentrations of all resources that have a species rate-limited by them. The steady state concentrations of the remaining resources can take any value as long as it is positive. Hence, in this limit the equations S2 can be viewed as a simple matrix test of whether a given set of surviving species limited by a given set of resources is possible for a given set of nutrient fluxes. Indeed, my multiplying the vector of fluxes with the inverse of the matrix 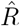 composed of inverse yields of surviving species and 1 for nutrients not limiting the growth of any species one formally gets the only possible set of steady state species abundances, 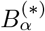, and a subset of non-limiting resource concentrations 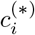 and 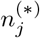. If all of them are strictly positive - the steady state is possible. If just one of them enters the negative territory - the steady state cannot be realized for these fluxes of nutrients.

The above rule can be modified to apply even below the high-flux limit with the following modifications: 1) Instead of *φ*^(*c*)^ (or *φ*^(*n*)^), one uses their “effective values” 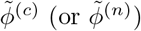 introduced in^19^, determined as

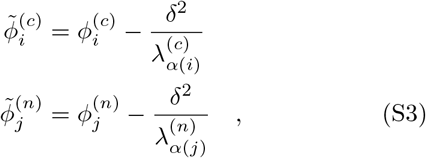

where *α*(*i*) is the (unique) species limited by the nutrient *i*. If the nutrient is not limiting for any os the species in the steady state, *α*(*i*) is the species using the nutrient in a non-limited fashion, which has the *smallest* value of λ. This last rule comes from the observation that in order for a non-limiting resource not to become limiting for a species *β* currently using it in a non-limiting fashion, its concentration cannot fall below 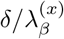. Thus, when checking the feasibility of a given state, the concentration of a non-limiting resource can be written as 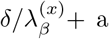 positive number, or (more conveniently) the influx of this resource can be offset as described in Eqs. S3

### Supplementary Note 3. Stable matching approach to identifying and counting uninvadable states

First we describe the exact one-to-one mapping between all uninvadable steady states (UIS) in our model and the complete set of “stable marriages” in a variant of a well-known stable marriage or stable allocation problem developed by Gale and Shapley in the 1960s^52^ and awarded the Nobel prize in economics in 2012. This mapping provides us with constructive algorithms to identify and count all uninvadable steady states in our ecosystem.

We start by considering a special case of our problem with *L* carbon and *L* nitrogen sources and a pool of *L*^2^ species, such that for every pair of sources *c*_*i*_ (carbon) and *n*_*j*_ (nitrogen) there is exactly one microbe *B*_*ij*_ capable of using them. For the sake of simplicity we have switched the notation from *B*_*α*_ to *B*_*ij*_, where *α* = (*ij*) is the unique microbe in our pool capable of growing on *c*_*i*_ and *n*_*j*_. Having considered this simpler situation we will return to the most general case of unequal numbers of carbon (*K*) and nitrogen (*M*) resources and any number of microbes from a pool of *S* species competing for a given pair of resources.

In what follows we will refer to a resource as *occupied* if in a given steady state there is a microbe for which this resource is rate-limiting. In our network representation occupied resources have an outgoing edge (their out-degree is equal to 1), while unoccupied resources have out-degree equal to 0.

#### Review of results about stable matchings in the hospitals/residents problem

The hospitals/residents problem^52^ is known in various settings. The one directly relevant to our problem is the following. There are *L* applicants for residency positions in *L* hospitals. A hospital number *i* has *L*_*i*_ vacancies for residents to fill, *L*_*i*_ ranging from zero to *L*. Each hospital has a list of preferences in which residency applicants are strictly ordered by their ranks, from 1 (the most desirable) to *L*, (the least desirable). These lists are generally different for different hospitals. Each applicant has a ranked list of preferred hospitals ranging from 1 (the most desirable) to *L* (the least desirable). Those lists can also vary between applicants. A *matching* is an assignment of applicants to hospitals such that all applicants got residency and all hospital vacancies are filled.

A matching is *unstable* if there is at least one applicant *a* and hospital *h* to which *a* is not assigned such that:

1. Condition 1. Applicant *a* prefers hospital *h* to his/her assigned hospital;
2. Condition 2. Hospital *h* prefers applicant *a* to at least one of its assigned applicants.

If such a pair (*a*, *h*) exists, it is called “a blocking pair” or “a pair that blocks the matching”. A *stable* matching by definition has no blocking pairs. Gale and Shapley proved that for any set of applicant/hospital rankings and hospital vacancies there is at least one stable matching^52^. Generally the number of stable matching is larger than one. For example, for stable marriages and random rankings the average number of stable matchings is given by *L*/*e* log *L*^53^. To the best of our knowledge, the dependence of this number on the distribution of hospital vacancies has not been investigated. The fact that the actual number of uninvadable states is rather close to its lower bound (compare black symbols and dashed line in Fig. 1) indicates that, at least for *L* ≤ 9, the number of stable matchings averaged over all possible in-degree allocations is rather close to 1.

Gale and Shapely not only proved the existence of at least one stable matching, but also proposed a constructive algorithm on how to find it. Listed below are the main steps in this algorithm optimized for for applicants. each applicants first submits his/her application to the hospital ranking 1 in his/her preference lists. Each hospital considers all applications it received so far and accepts all of the applicants if their number is less or equal than hospital’s announced number of vacancies, *L*_*i*_. If the number of applicants exceeds *L*_*i*_, the hospital gives a conditional admission to the best-ranking *L*_*i*_ applicants according to hospital’s own preference list. Each applicant not admitted to their top hospital goes a step down on his/her preference list and applies to the second-best hospital. The latter admits this applicant if (1) this hospital has not yet filled all of its vacancies or (2) all vacancies are filled, but among the conditionally admitted applicants there is at least one who ranks lower (according to hospital’s list) than the new applicant. Such lower-ranked applicants are declined admission and replaced with better ones. They subsequently lower their expectations and apply to the next hospital on their list. After a number of iterations all applicants are admitted and all vacancies are filled so that this process stops. As Gale and Shapley proved in Ref. ^52^, the resulting matching is stable. Furthermore, the theorem states that in this matching every applicant gets admitted to the best hospital among all stable matchings, while every hospital gets the worst set of residents among all stable matchings. Later research described in Ref.^53^ describe more complex constructive algorithms allowing one to efficiently find all of the stable matchings starting with the applicant-optimal one.

Well developed mathematical apparatus of stable matching problem provides an invaluable help in the task of identifying all uninvadable states in microbial ecosystems. Indeed, without its assistance this task would require exponentially longer time. To connect the problem of finding all uninvadable states to that of finding all stable matchings between hospitals and residents, we start with the following three observations:

1. In any uninvadable steady state, either all carbon sources or all nitrogen sources (or both) are occupied. Indeed, if in a steady state a carbon source *c*_*i*_ and a nitrogen source *n*_*j*_ are not-limiting to any microbes, then microbe *B*_*ij*_ can always grow and thereby invade this state. Thus uninvadable states can be counted separately: one first counts the states where all nitrogen sources are occupied, and then counts those in which all carbon sources are occupied. Double counting happens when both carbon and all nitrogen sources are occupied. We will keep the possibility of double counting in mind and return to this problem later.
2. For a pool of species, where for every pair of resources there is exactly one microbe using each carbon and each nitrogen. One can think of each of *L* carbon (alternatively, nitrogen) sources as if it has a list of “preferences” ranking all nitrogen (correspondingly carbon) sources. Indeed, the ranking of competitive abilities 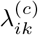 of different microbes using the same carbon source *c*_*i*_ but different nitrogen sources *n*_*k*_ can be viewed as the ranking of nitrogen sources *k* by the carbon source *i*. Conversely, the ranking of 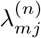 with the same *n*_*j*_ but variable *c*_*m*_ can be thought of as ranking of carbon sources *c*_*m*_ by the nitrogen source *n*_*j*_.
3. Consider a steady state in which all nitrogen sources are occupied. In our network representation it corresponds to every nitrogen source sending an outgoing link to some carbon source. Let *L*_*i*_ be the number of microbes using the carbon source *i* in a non-limiting fashion (the in-degree of these outgoing links ending on *c*_i_). Then, obviously, *L* = ∑ *L*_*i*_ (note that some of the terms in this sum might be equal to zero).

One can prove that if the state is uninvadable, then the matching given by all edges going from nitrogen sources to carbon sources must be stable in the Gale-Shapley sense. To prove this, let’s think of nitrogen sources as “applicants” and nitrogen sources as “hospitals” with their numbers of “vacancies” given by *L*_*i*_. Indeed, any unstable matching has at least one blocking pair (*n*_*j*_, *c*_*i*_) such that:

- Condition 1. The nitrogen source (‘applicant”) *n*_*j*_ “prefers” the carbon source (“hospital”) *c*_*i*_ to its currently assigned carbon source (the one used by the current microbe *B*_*kj*_ limited *n*_*j*_). This means that 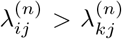 Thus the microbe *B*_*ij*_ can grow on its nitrogen source (provided that it can also grow on its carbon source).
- Condition 2. The carbon source (“hospital”) *c*_*i*_ “prefers” the nitrogen source (“applicant”) *n*_*j*_ to at least one of *L*_*i*_ of its currently assigned carbon sources (the set of microbes using *c*_*i*_ in a non-rate-limiting fashion). Thereby 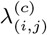 must be larger than the smallest λ^(*c*)^ among these microbes. According to the Exclusion Rule 2, this smallest λ^(*c*)^ is still larger than λ^(*c*)^ of the microbe limited by *c*_*i*_ (if it exists). Thus the microbe *B*_*ij*_ can also grow on its carbon source.

This proves that the microbe *B*_*ij*_ corresponding to any blocking pair can grow on both its carbon and its nitrogen sources, and thereby can successfully invade the steady state. This finishes the proof that any uninvadable state has to be a stable matching in the Gale-Shapley sense.

However, this does not prove that any stable matching corresponds to exactly one uninvadable state. To prove this we first notice that, up to this point, our candidate uninvadable state contained only the nitrogen-limited species. We will now supplement it with carbon-limited species in such a way that 1) added species do not violate the exclusion rule 2; 2) added species render the state completely uninvadable. Let is introduce a new notation (applicable to our case in which all nitrogen sources are occupied). Let 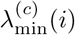 to denote the smallest λ^(*c*)^ among all species using *c*_*i*_ in a non-rate-limiting fashion. The Gale-Shapley theorem only guarantees the protection of our state from invasion by a species (*i*, *j*) with 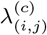 larger than 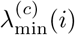 (see the Condition 2 above). To ensure that our state is uninvadable by the rest of the species, one needs to add some carbon-limited species to this state. In order to do this in a systematic way, for each *c*_*i*_ we compile the list of all species using this carbon source with 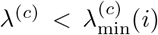. Each of these species is a potential invader. Some species could be crossed off from the list of potential invaders because they cannot grow on their nitrogen source. These species have λ^(*n*)^ below that of the (unique) species limited by their nitrogen source. Among the species that remained on the list of invaders after this procedure, we select that with the largest λ^(*c*)^ and add it to our steady state as a *C* → *N* directed edge, that is to say, as a carbon-limited species. This will prevent all other potential invaders on our list, since they have smaller λ_(*c*)_ and thus, following the addition of our top carbon-limited species, they would no longer be able to grow based on their carbon source. We will go over all *c*_*i*_ and add such carbon-limited species if they are needed. The only scenario when such species is not needed if our list of potential invaders would turn up to be empty. In this case we will leave this carbon source unoccupied. Since for each carbon source the above algorithm selects the carbon-limited species (or selects to add no such species) in a unique fashion, there is a single uninvadable state for every stable matching in the Gale-Shapley sense. We are now in a position to predict and enumerate all uninvadable states in our model.

#### Lower bound on the number of uninvadable states

To count the number of partitions (*L*_1_, *L*_2_,…, *L*_l_) such that ∑ *L*_*i*_ = *L*, one can use a well known combinatorial method. According to this method, one introduces *L* − 1 identical “separators” (marked with |) which are placed between *L* identical objects (marked ·) separating them into *L* (possibly empty) partitions. For example, for *L* = 4 a partition 0, 1, 0, 3 would be denoted as | · || …. The combinatorial number of all possible arrangements of separators and objects is obviously 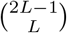. For every such partition the Gale-Shapley theorem guarantees at least one stable matching (that is, at least one uninvadable steady state). The lower bound on the number of uninvadable steady states has to be doubled to account for reversal of roles of carbons and nitrogens. There is a small possibility that we double counted one partition (1, 1, …, 1). Indeed, the unique uninvadable stable state corresponding to this partition could in principle be counted both when we start from nitrogen sources and when we start from carbon sources. This could happen only when the numbers of carbon and nitrogen sources are equal to each other. More restrictively, this partition will be double-counted only if, when we started from C, all of the N-sources will send a link back to C, and these links all will end on different C-sources. The same has to be true if one starts with N-sources and at then sends links back to C. The steady state network in this case will consist of one or more loops covering all nutrients. However, one can prove that, at least for the Gale-Shapley nitrogen-optimal state, the last carbon to be picked up would not need to send back a carbon-limited link. Thus in our task of calculating the lower bound on the number of uninvadable states, we don’t need to correct for the possibility of double-counting since at least one stable matching per partition (namely the Gale-Shapley) would not be double-counted. Then we have 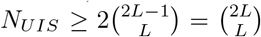 The Sterling approximation for this expression is 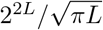. Thus the overall lower bound for the number of uninvadable stable states is given by

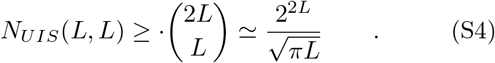

More generally, the number of carbon sources, *K*, is not equal to the number of nitrogen sources, *M*. The resource type with a larger number will always have at least one resource left without input. Thus here one never needs to correct for double counting. Using the same reasoning as for *K* = *M* = *L*, the lower bound on the number of resources in this case is given by 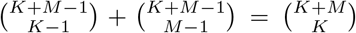 Here, the first term counts the uninvadable steady states in which all nitrogen sources are occupied and the partition divides *M* edges sent by nitrogen sources among *K* carbon sources, which requires *K* − 1 “dividers”. The second term counts the number of uninvadable steady states in which all carbon sources are occupied. Denoting the fraction of carbon resources among all resources as *p* = *K*/(*K* + *M*) and using the Stirling approximation one gets

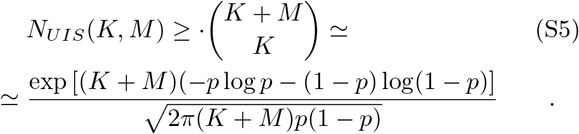

In the case of multiple microbial species using the same pairs of resources, our version of the Gale-Shapley resident-oriented algorithm must be further updated. Let *M* be the number of nitrogen sources, and *K* — the number of carbon sources in the ecosystem, *S* the number of species in our pool, each requiring a pair of resources to grow. As now there may be more than one microbe that uses a given pair of resources *c*_*i*_ and *n*_*j*_, we introduce the notation 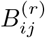 for the *r*th microbe using the same pair of sources *c*_*i*_ and *n*_*j*_. On average, each nitrogen (carbon) source has *S*/*K* (*S*/*M*) microbes, which are capable of using it. As in the traditional Gale-Shapley algorithm, each nitrogen (carbon) source ranks all microbes capable of using it by their λ^(*n*)^ (λ^(*c*)^).

The way to identify all uninvadable stable states in this case is determined by a variant of the stable marriage problem (or rather the hospital/resident problem) in which every man (and every woman) may have more than one way to propose marriage to the same woman (man). In our model this corresponds to more than one microbe (a type of marriage) capable of growing on the same pair of carbon (corresponding to, say, men) and nitrogen (corresponding to women) sources. You may think of it as if each participant has several different ways to propose to the person of the opposite sex (send flowers, take to a restaurant, etc). Each of these proposals is ranked by both parties independent of other ways. As far as we know, this variant has not been considered in the literature yet. However, all of the results of the usual stable marriage (or hospital-resident) problem remain unchanged.

One can easily see that our lower bound (Eq. S5) on the number of uninvadable states (equal to the number of stable marriages in all partitions) remains unchanged. Indeed, it is given by the number of partitions and hence depends only on *K* and *M* and not on *S*. However, for *S* ≫ *K* · *M* one expects to have many more stable marriages for each partition. Thus the lower bound we have established is likely to severely underestimate the actual number of UIS in the ecosystem. Indeed, according to the SI section “The number of uninvadable states in a continuous approximation”, the number of uninvadable states grows much faster than the lower bound of *O*(2^*K*+*M*^). Future work is needed to connect the stable marriage results to those derived in the continuous approximation (see Eq. S28 below).

### Supplementary Note 4. The number of allowed states and the number of uninvadable states in a continuous approximation

: We can calculate the number of allowed and, separately, uninvadable states in our model in the limit of *K*, *M* ≫ 1 and *S* ≫ *K*, *M*. In this limit, every nutrient has a large average number of microbes competing for its utilization. Throughout this section we assume that each of the resources has equal number of microbes capable of using it (*S*/*K* for carbon and *S*/*M* for nitrogen). Let 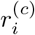 (respectively 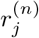) be the rank of the (unique if present) microbe whose growth is limited by *c*_*i*_ (respectively *n*_*j*_). The rank is defined as the the number of microbes in the pool with value of λ larger or equal than *r*. Hence, in our pool of species, the most competitive microbe for each nutrient has the rank 1, while the worst one - the rank *S*/*K* (or *S*/*M* for nitrogen resources). It is convenient to assume that in a special case, where there is no microbe limited by the resource, the rank of the resource is equal to *S*/*K* + 1 (*S*/*M* + 1 correspondingly). In this case all microbes using this nutrient would be allowed to grow on it according to our competitive exclusion rules. It is also convenient to normalize the ranks as 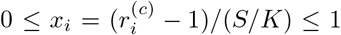 and 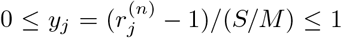. These normalized variables quantify the probability that a randomly selected microbe using a given nutrient (*c*_*i*_ for *x*_*i*_ and *n*_*j*_ for *y*_*j*_) would be able to grow on it (provided that the second resource would also allow for its growth). To calculate the probability *x* that a randomly selected microbe would be able to grow on its carbon source one has to average *x*_*i*_ over all carbon sources: *x* = ∑_*i*_ *x*_*i*_/*K*. Similarly, the probability for a random microbe to be able to grow on its nitrogen source is given by *y* = ∑_*j*_ *y*_*j*_/*M*. In what follows we will carry out the summation over all possible values of all normalized ranks of carbon, *x*_*i*_ (0: 1/(*S*/*K*): 1), and nitrogen, *y*_*j*_ (0: 1/(*S*/*M*): 1), sources. In the continuous limit *K*/*S*, *M*/*S* ≪ 1 these sums can be replaced by integrals over continuous variables ranging between 0 and 1. For *K* ≫ 1, the average rank *x* of all carbon sources has an approximately Gaussian distribution with width 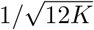, while the average rank *y* of all nitrogen sources has a Gaussian distribution with width 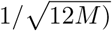. Indeed, the variance of the uniform distribution between 0 and 1 is 1/12, while the variance of the average is reduced by the number of variables in the sample. Some of our calculations require knowledge of the probability density function outside the region of validity of central limit theory. We have also carried out calculations using the exact PDF of the sum of *K* (or *M* in case of nitrogen) uniformly distributed variables known as Bates distribution (see Eq. (2) in http://mathworld.wolfram.com/UniformSumDistribution.html. for the exact functional form of the PDF of the Bates distribution). The results for the Bates distribution were very close to those for the Gaussian distribution. To account for significant difference at *K* = *M* = 1, Fig. 1 shows our calculations using the Bates distribution (red and black long-dashed lines). Hence, in what follows we will consider only the Gaussian case.

#### The number of allowed states in a continuous approximation

Let us first calculate the number of allowed states. Consider a state in which 0 ≤ *K*_*L*_ ≤ *K* of carbon sources and 0 ≤ *M*_*L*_ ≤ *M* of nitrogen sources each have (a unique) microbe limited by them (and making them limited). The combinatorial number of choices of such microbe-limited resources is given by 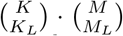. The number of ways to choose one limiting microbe on each of these resources is given by *S*/*K* for carbon resources and *S*/*M* for nitrogen resources. Indeed, since each species has exactly one carbon (nitrogen) source it could utilize, the number of species per each resource is *S*/*K* (*S*/*M* correspondingly). The total number of ways to choose *K*_*L*_ + *M*_*L*_ microbes in the candidate steady state is thus given by 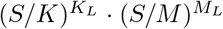. The probability that all these microbes would be allowed by their non-limiting resources is given by 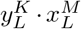 (note that the average rank *y* of nitrogen resources is raised to the power of *K*_*L*_ of limited carbon sources and vice versa). Indeed, the selection of a non-limiting resource is entirely random when one goes over all possible microbe candidates. The sum over all possible values *K*_*L*_ and *M*_*L*_ is simply given by:

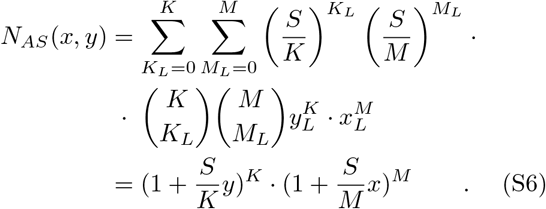

Thus the total number of the allowed states (invadable or not) is given by the following integral:

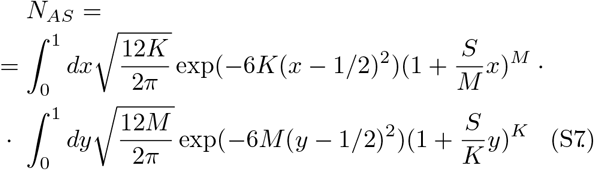

This integral can be calculated using the saddle point approximation. In the limit *M* ∼ *K* ≫ 1 and *M*/*S* ≫ 1, the saddle point *x*\* for the first integral over *x* is given by

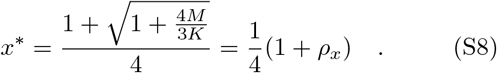

Here in order to simplify the notation we introduced a new variable 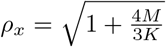. The integral over *x* in the saddle point approximation is then given by

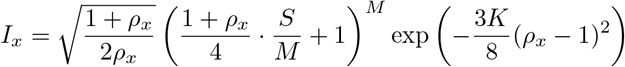

Similarly, the integral over *y* in the saddle point approximation is then given by

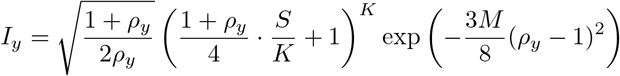

Here 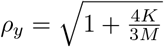 and is related to *ρ*_x_ by

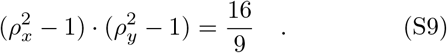

The number of allowed steady states *N*_*AS*_ is simply the product of *I*_*x*_ and *I*_*y*_. In the symmetric limit of the equal number of nutrient sources *M* = *K* = *L* where 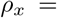 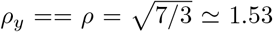 the formula for the number of steady state can be simplified as

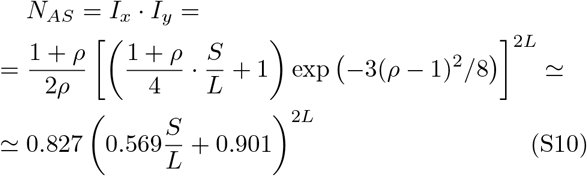

As one can see the number of allowed states rapidly increases with both the number of resources of each type, *L*, as well as with number of species per each resource, *S*/*L*. This increase however is much slower than that in the number of candidate states not constrained by the exclusion rule 2. Indeed, the number of such candidates *N*_*c*_ = (1 + *S*/*K*)^*K*^ · (1 + *S*/*M*)^*M*^, which for *K* = *M* = *L* becomes (*S*/*L* + 1)^2*L*^ (compare this expression to Eq. S10).

Finally, for *S* = *L*^2^ used in our simulations shown in Fig. 1, the expression for the number of allowed states becomes *N*_*AS*_ ≃ 0.827 (0.569*L* + 0.901)^2**L**^.

#### The number of uninvadable states in a continuous approximation

To calculate the number of uninvadable states, one needs to check if each of the allowed states calculated in the Eq. S7 can be invaded by each of the species that are currently not present in the state. Fortunately, the notation introduced in the previous section makes this task very easy. When calculating the number of allowed states in our model we were going over all species present in the state and multiplying our formula by the probability that it’s *non-limiting* resource is allowed by our rules of competitive exclusion. This probability is equal to *y* = ∑_*j*_ *y*_*j*_/*M* for species limited by the concentration of their carbon sources and *x* = ∑_*i*_ *x*_*i*_/*K*. Here (as before) *x*_*i*_ and *y*_*j*_ are the normalized ranks of the species limited by *c*_*i*_ and *n*_*j*_ correspondingly. *x*_*i*_ = 0 (or *y*_*j*_ = 0) corresponds to a situation where there are no species in our pool with λ larger then the species currently limited by this carbon (or nitrogen) source. Conversely, *x*_*i*_ = 1 corresponds to a situation where the resource is currently not limiting for any of the species in the steady state under consideration. Hence the growth of any introduced species on this resource is allowed by the competitive exclusion rules. *x*_*i*_ = 1/2 corresponds to a case where the species with a median value of λ^*c*^ is limiting *c*_*i*_ so that exactly half of all species using this resource can grow on it (provided that their nitrogen source allows for growth). A species not currently present in the steady state can grow in it if and only if *both* its carbon and nitrogen sources allow for its growth. The probability of this being true for a randomly selected species is simply *x* · *y*, while the probability that it is not allowed to grow by either one or both of its nutrients is simply 1 − *x* · *y*. The probability that none among *S* species can grow in a given steady state is

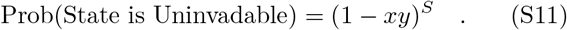

Here in principle we only need to check uninvadability against *S* − *K*_*L*_ − *M*_*L*_ species *not present* in the steady state tested for invadability However, in the limit *S* ≪ *K*, *M*, this difference is small and would be ignored in our calculations.

Thus the number of uninvadable states is given by an integral similar to Eq. S7:

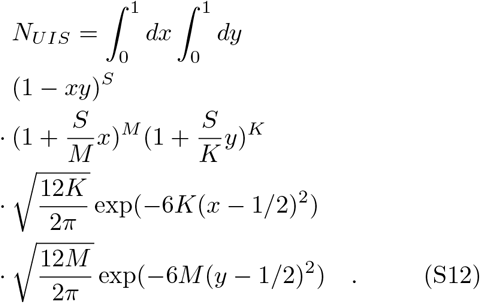

The density of stable states on the *x* − *y* plane described by this equation can be visualized already *L* = 9. For tables of λ^(*c*)^ and λ^(*n*)^ used in our main text we exhaustively identified 81,004 UIS. For each of these UIS we calculated *x*_*i*_ and *y*_*i*_ - the average normalized ranks of microbes limited by their carbon and nitrogen sources respectively. The natural logarithm, of the density of these 81,004 points on the *x* − *y* plane is visualized in Fig. S2. One can see that most states are localized within a “smile” stretching from the upper left corner (the top competitors for carbon and the weakest competitors for nitrogen) to the lover right corner (the top competitors for nitrogen and the weakest competitors for carbon) of the diagram. That means that the average ranks of carbon and nitrogen competitive abilities of microbes present in steady states and limited by these resources are negatively correlated.

**FIG. S2.**
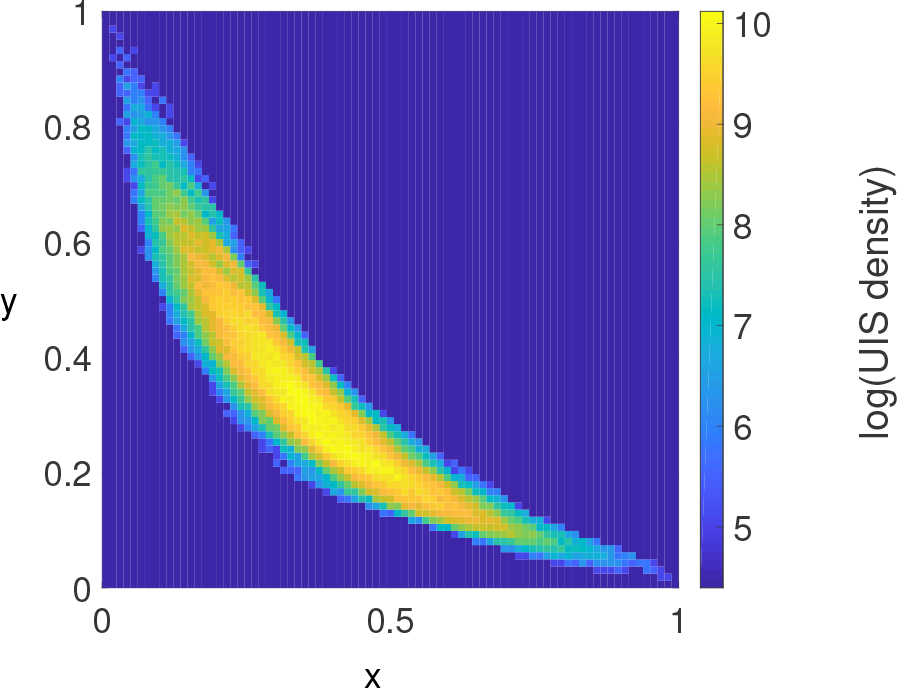
The density of UIS for *L* = 9. The pseudocolor plot shows the natural logarithm of the density of uninvadable stable states on the *x* − *y* plane, where *x* and *y* are, respectively, the average normalized ranks of carbon- and nitrogen-limited microbes across all resources.

The probability of the state being uninvadable couples the integration over *x* and *y* in the Eq. S12. This integral, when written as 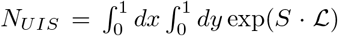, in the limit of large *S* ≫ *K*, *M* can be approximately calculated in the saddle point approximation.Here

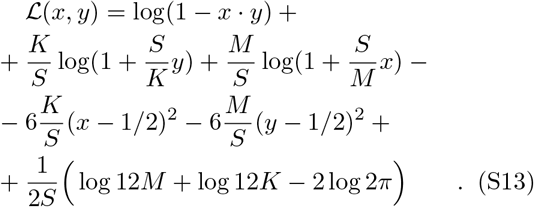

The integral of Eq. S12 over *x* is dominated by the saddle point *x*\*(*y*) obtained by solving for x the following equation

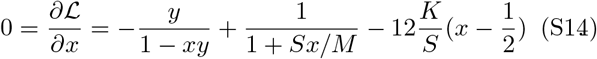

We will be interested in solving this equation in the regime where *x* ≃ 1 (hence 1/(1 + * *S*_*x*_/*M*) ≃ *M*/(*S*_*x*_)), and *y* ≫ 1 (hence − *y*/(1 − *xy*) ≃ −*y*. In this limit one gets a quadratic equation for *x*

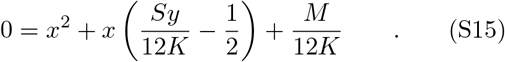

The solution is given by

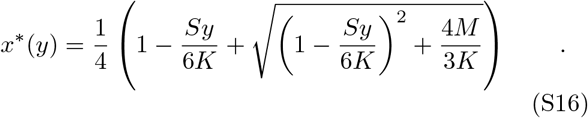

For *M* = *K* the maximal value of *x*\* (*y*) is reached at *y* = 0 and is equal to 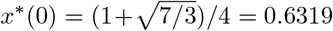. For large excess of the number of nitrogen sources over that of carbon ones, *M* > 6 *K*, *x*\*(*y*) can reach its maximal value of 1 even before *y* hits 0. In this case, the saddle point disappears and the integral will be dominated by region near *x* = 1. We will leave for future studies the calculation of the number of uninvadable steady states in this case. In the limit *M*/*S* ≪ 1, *K*/*S* ≪ 1 the second derivative evaluated at this stable point is

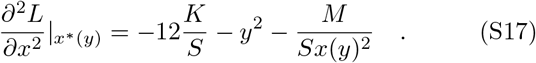

The saddle point integration over *x* results in the following expression for the number of uninvadable states

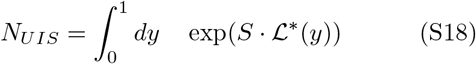

where

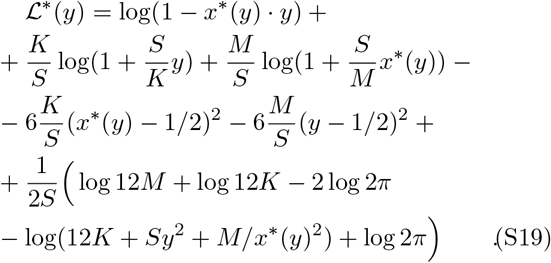

Here the last term 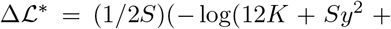 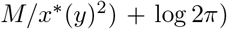 comes from the saddle point integral over *x*. In other words, 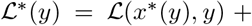 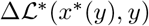, where 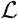 is defined by the Eq. S13. The integral over *y* has two saddle points: one for small *y* ∼ *K*/*S* and hence large *x*(*y*) ∼ 1 and the other for large *y* ∼ 1 and small *x*(*y*) ∼ *M*/*S*. We will calculate only the first saddle point and then apply symmetry arguments to extend these results to the second one. Indeed, if instead of integrating out *x* we were to integrate out *y* first, the order of saddle points will change places. Hence, we are interested only in the region of small *y* ∼ *K*/*S*. The saddle point is determined by 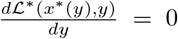. Let’s first calculate the derivative of the last term (referred to as 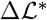) in the Eq. S19. It is given by 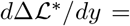 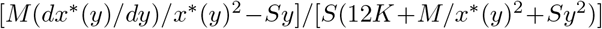. The first term in the enumerator and the first two terms in the denominator dominate the expression for *y* ∼ *K*/*S* resulting in

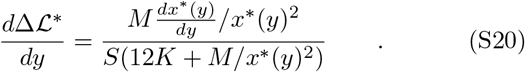

While *dx*\*(*y*)/*dy* ∼ *S*/*K* is large, the whole expression is still of order of 1/*M* or 1/*K* and hence much smaller than 1. As we will see below, the dominant contribution to 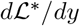 is of order of 1. Hence, 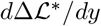 can be safely ignored. One then has 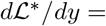 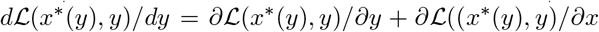. *dx*\*/*dy*. Since the saddle point integration over *x* required that 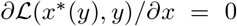, the second term is zero. The first term is given by 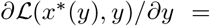 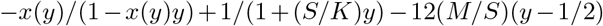. In the limit *y* ∼ *K*/*S* and *x*(*y*) = *O*(1), the first two terms are of order of 1, while the third term can be ignored. Furthermore, the denominator in the first term can be ignored. Hence, the saddle point *y*\* is determined by the following condition

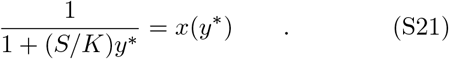

By introducing the dimensionless variable 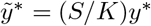 and plugging it into the Eq. S16 one gets

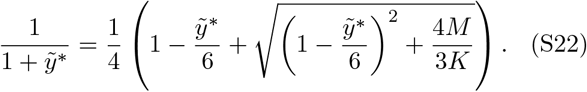

While in general this equation does not have the analytic solution, it can be easily solved numerically for any value of *M*/*K* < 6 (the saddle point disappears for *M* > 6*K*). To fit our numerical simulations of the model with *M* = *K* = *L* and *S* = *L*^2^, we solved Eq. S22 for *M*/*K* = 1:

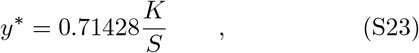

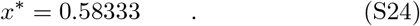

Note that both *x*\* and *y*\* are far away from 1/2 and hence are located where the Gaussian approximation to the sum of uniformly-distributed random numbers no longer applies. However, the Gaussian was not among the main factors deciding the position of the fixed point. Thus, the results derived above could still be used. We confirmed it by carrying the saddle point calculations in Eq. S18 using the exact form of the Bates distribution describing the sum of *M* uniformly-distributed random numbers. Much as for the number of allowed states, for large *L* the number of uninvadable states calculated using the Bates distribution was very close to the same number calculated using the Gaussian distribution.

To further verify the accuracy of our calculations of *x*\* and 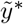 in Eqs. S24 and S23 correspondingly, in Fig. ?? we plotted 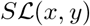 as a function of *x* and *y* for *K* = *M* = *L* and *S* = *L*^2^. The red dot marking the predicted position of the saddle point is in excellent agreement with its numerically-determined position.

Hence, for large *L* most uninvadable states come from two regions on the *x* − *y* plane. Let us briefly return to the original notation for which the rank of the most competitive microbe using each nutrient is 1, while that of the least competitive one is *S*/*K* for carbon sources and *S*/*M* for nitrogen sources. Uninvadable states contributing to the saddle point above must have many microbes near the top of the nitrogen ranking table with average nitrogen rank being 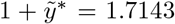. This can be realized e.g. if around 71% of microbes were the second best competitors for their nitrogen resource, while about 29% were its best competitors In the whole pool. There are of course many other solutions all giving the average ranking shown above. On the opposite side, the average rank of microbes on their carbon resources is in the middle of the table *x*\* (*S*/*K*) = 0.5833*L*. The other saddle point corresponds to carbon and and nitrogen swapping places with each other.

**FIG. S3.**
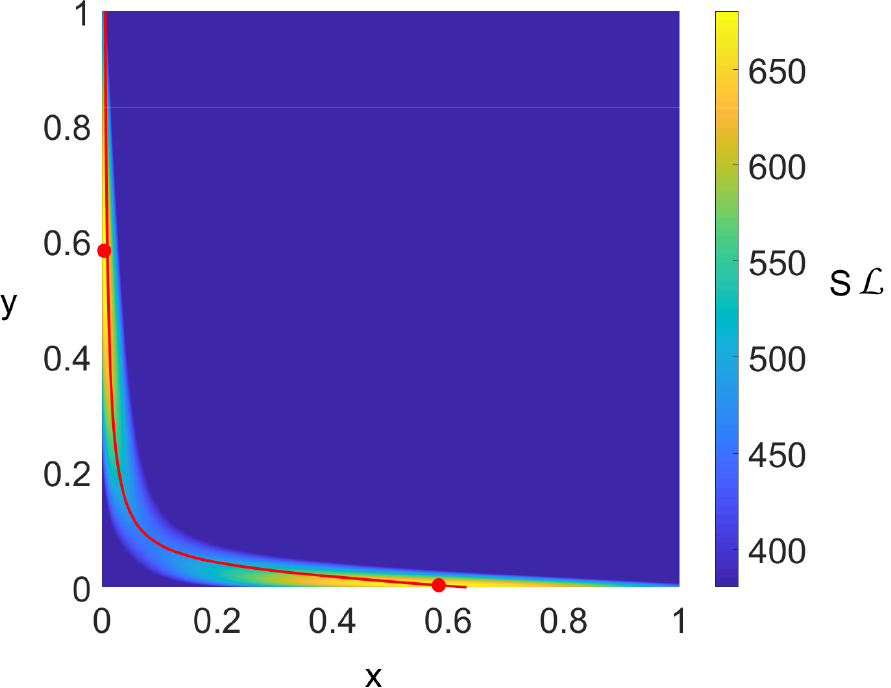
The logarithm of the density of UIS for *L* = 200 along with the saddle point line. *x*\*(*y*) **vs** *y*. The pseudocolor plot shows 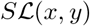 - the logarithm of the density of UIS on the *x* − *y* plane calculated for *L* = 200 as a function of normalized carbon and nitrogen average ranks *x* and *y*. The red line follows *x*\*(*y*) vs *y* described by the Eq.S16. The red dots mark the predicted positions of two saddle points according to Eqs. S24 and S23 and a symmetric one with *x* and *y* swapped places.

The second derivative 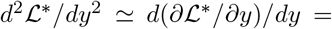 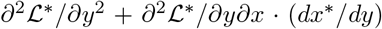. Here again we ignored 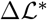 since its contribution to the derivative is 1/*K* smaller than the above terms. The first term is given by 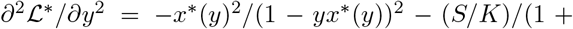 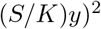. Here the second term is much larger. One also has 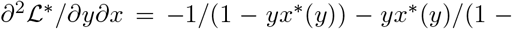 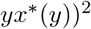. Here the first term is much larger. Given that *dx*\*(*y*)/*dy* ∼ *S*/*K*, the dominant contributions to 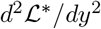 from 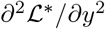 and 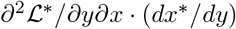 are comparable to each other and are both of order of *S*/*K*. The derivative *dx*\*/*dy* is given by

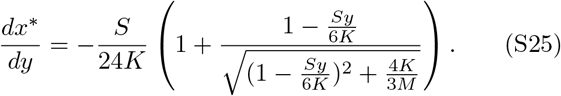

Evaluating this expression for *K* = *M* at *y*\* = 0:71428*K*/*S* one gets 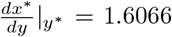 Hence, the final expression for 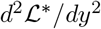 at the saddle point is

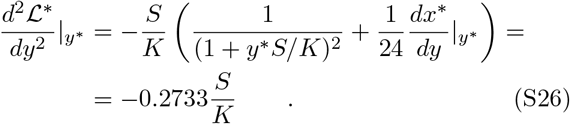

**FIG. S4.**
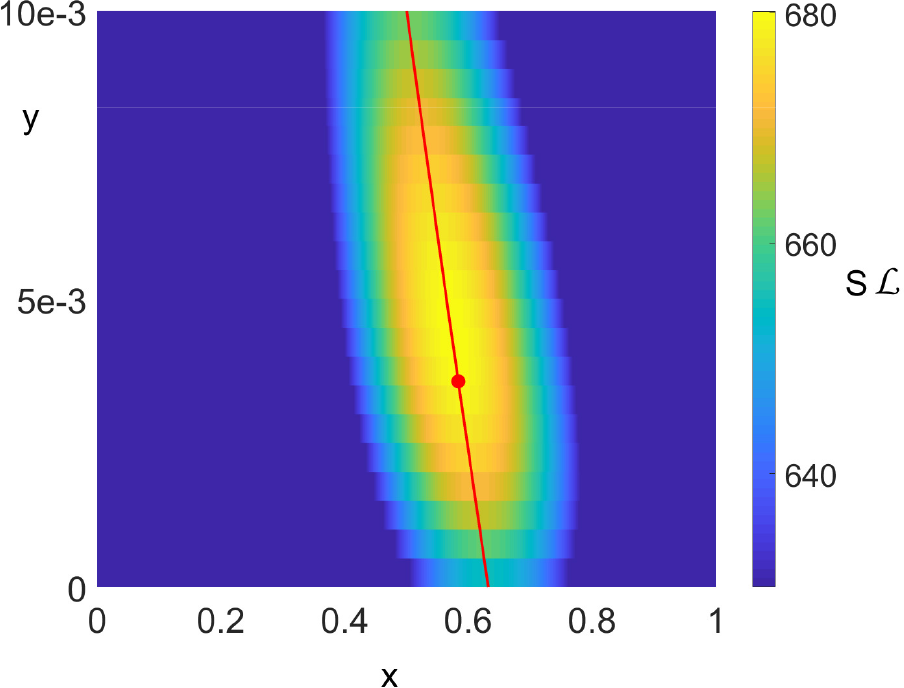
The logarithm of the density of UIS for *L* = 200 along with the saddle point position. The pseudocolor plot shows 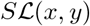 - the logarithm of the den-sity of UIS on the *x* − *y* plane calculated for *L* = 200 as a function of normalized carbon and nitrogen average ranks *x* and *y*. The red dot marks the predicted position of the saddle point according to Eqs. S24 and S23. It is in excellent agreement with its numerically-determined position.

**FIG. S5.**
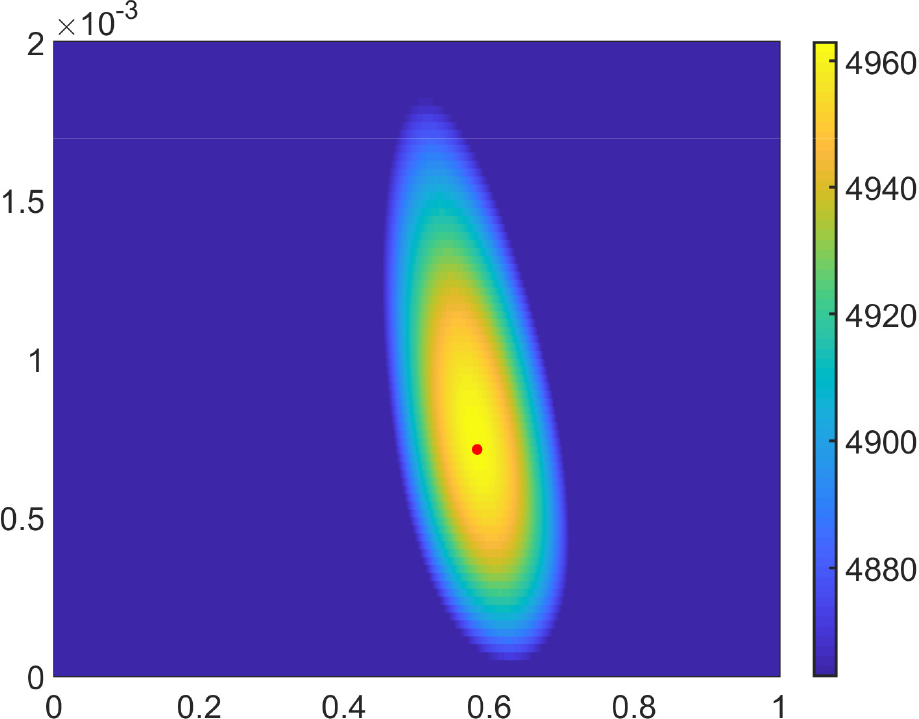
The density of UIS for *L* = 1000 along with the saddle point position. The pseudocolor plot shows 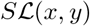 calculated for *L* = 1000 as a function of *x* and *y*. The red dot marking the predicted position of the saddle point
according to Eqs. S24 and S23 is in excellent agreement with its numerically-determined position.

Plugging Eqs. S23, S24, S26 into the saddle point estimate of the integral given by Eqs. S18, S19 one gets (term-by-term):

- 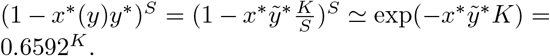
- 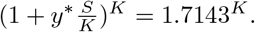
- 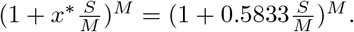
- 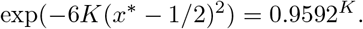
- 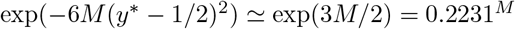
- The saddle point integration over *x* combined with the normalization constant of the Gaussian distribution of *x* generates

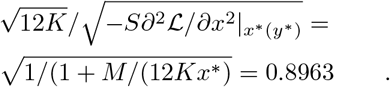
- The saddle point integration over *y* combined with the normalization constant of the Gaussian distribution of *y* generates

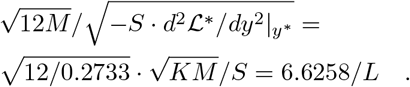
- Factor 2 for *K* = *M* takes into account that our calculations were done for one saddle point in which 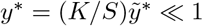 and *x*\* = *O*(1). The symmetric point located diagonally across this one on the *x*−*y* plane would have an identical contribution. For *K* ≠ *M* one of these saddle points would dominate the asymptotic formula.

Once all these terms are put together one gets the following asymptotic formula for the number of uninvadable steady states for *L* ≫ 1, and *K* = *M* = *L*.
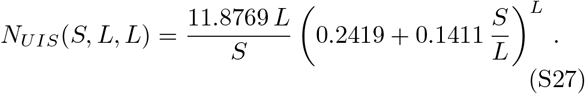

If in addition, the number of species in the pool is given by *S* = *L*^2^ (as used in our numerical simulations for small *L*), one gets

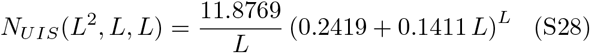

For *K* ≠ *M* this formula needs to be modified by first solving the Eq. S22 to find the new saddle points *x*\* and *y*\*. These values then need to be plugged into the bullet list shown above to update the numerical coefficients combined in Eq. S27. One fact remains generally true, however. The leading super-exponential contribution would be given by

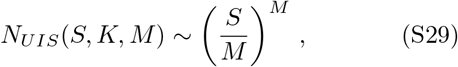

where *M* corresponds to the resource with the largest number of nutrients (nitrogen in this example, where we assumed that *M* > *K*).

To test how well this expression calculated in the limit *L* ≫ 1 matches the numerical integration of Eq. S12, in Fig. S6 we compare them for *L* going up to 1000 (and *S* = *L*^2^). Fig. S6 plots the number of uninvadable states raised to the power of 1/*L* plotted as a function of *L*. The black symbols correspond to the 2-dimensional numerical integration of Eq. S12, where both *x* and *y* range between 0 and 1 in steps of 1/*L*^2^. Because of numerical limitations, the integration has been only carried for *L* ≤ 100. The red line is the 1-dimensional numerical integration of the Eq. S18 over *y*. Now *L* is extended up to 1000. The blue line is given by the complete saddle point calculation (Eq. S28). The ratios between either two of these three expressions asymptotically converges to 1. Note that the saddle point expression in Eq. S28 is completely off for *L* ≤ 9 shown in Fig. 1.

**FIG. S6.**
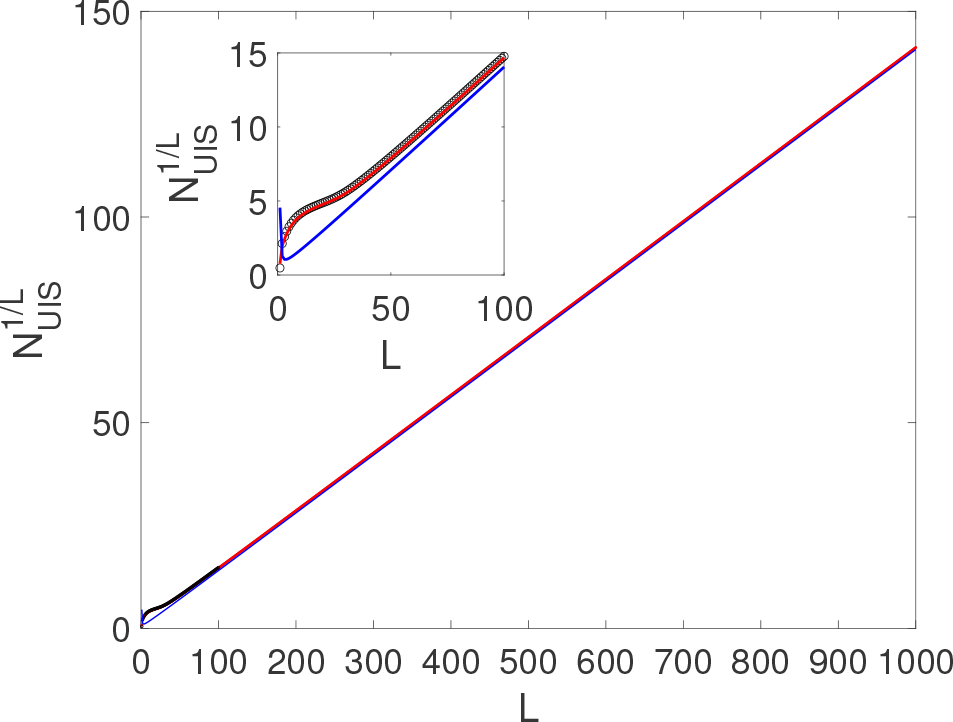
Saddle point approximation to the number of UIS in the continuous limit. The number of uninvadable states raised to the power of 1/*L* plotted as a function of *L*. The black symbols correspond to the 2-dimensional numerical integration of Eq. S12, where both *x* and *y* range between 0 position of the saddle point according to Eqs. S24 and S23 is in excellent agreement with its numerically-determined position. Plugging Eqs. S23, S24, S26 into the saddle point estimate of the integral given by Eqs. S18, S19 one gets (term-by-term): and 1 in steps of 1/*L*^2^. Because of numerical limitations, the integration has been only carried for *L* ≤ 100. The red line is the 1-dimensional numerical integration of the Eq. S18 over *y*. Now *L* is extended up to 1000. The blue line is given by the complete saddle point calculation (Eq. S28). The ratios between either two of these three expressions asymptotically converges to 1. The inset zooms up on the region 1 ≤ *L* ≤ 100.

#### The number of allowed and uninvadable states for more than two types of essential resources: continuous representation

Note that all of the above formulas could be easily gen-eralized to a biologically meaningful case of more than two types of essential nutrients. For example, if one was to add another essential nutrient type (e.g.sources of phosphorus), the normalized ranking of λ^(*P*)^ would introduce a new variable 0 ≤ *z* < 1. The above formulas would be modified so that the probability that a state is not invadable now becomes

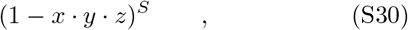

while the combinatorial factor calculating the number of allowed states for a given value of the average ranks *x*, *y*, and *z* is given by

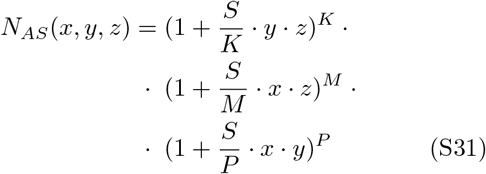

Here *P* is the number of sources of phosphorous in the system, and the equation above sums over all microbes that are limited by either C, N, or P. It takes into account that a microbe limited by one resource has to be able to grow on the other two resources.

### SUPPLEMENTARY TABLES

**TABLE I.** 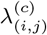, 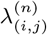 values of the 4 species for the 2C×2N×4S model.

| | $\lambda_{(i,j)}^{(c)}$ | | $\lambda_{(i,j)}^{(n)}$ | |
| --- | --- | --- | --- | --- |
| | $N_1$ | $N_2$ | $N_1$ | $N_2$ |
| $C_1$ | 41 | 35 | 16 | 50 |
| $C_2$ | 52 | 56 | 27 | 44 |

**TABLE II.** Values of carbon and nitrogen Yields of the 4 species for the 2C×2N×4S model.

| | $Y_{(i,j)}^{(c)}$ | | $Y_{(i,j)}^{(n)}$ | |
| --- | --- | --- | --- | --- |
| | $N_1$ | $N_2$ | $N_1$ | $N_2$ |
| $C_1$ | 0.37 | 0.64 | 0.35 | 0.50 |
| $C_2$ | 0.47 | 0.14 | 0.40 | 0.30 |

**TABLE III.** 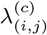 values of the 36 species for the 6C×6N×36S model.

| | $N_1$ | $N_2$ | $N_3$ | $N_4$ | $N_5$ | $N_6$ |
| --- | --- | --- | --- | --- | --- | --- |
| $C_1$ | 47.4 | 78.1 | 93.7 | 68.9 | 75.0 | 44.5 |
| $C_2$ | 89.6 | 68.6 | 33.6 | 77.8 | 16.5 | 90.8 |
| $C_3$ | 56.5 | 32.2 | 86.2 | 13.1 | 71.1 | 15.5 |
| $C_4$ | 53.0 | 94.1 | 38.7 | 10.7 | 34.0 | 34.9 |
| $C_5$ | 25.0 | 49.3 | 76.3 | 18.2 | 54.5 | 51.8 |
| $C_6$ | 47.1 | 91.9 | 57.7 | 63.0 | 92.2 | 90.0 |

**TABLE IV.** 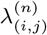 values of the 36 species for the 6C×6N×36S model.

| | $N_1$ | $N_2$ | $N_3$ | $N_4$ | $N_5$ | $N_6$ |
| --- | --- | --- | --- | --- | --- | --- |
| $C_1$ | 18.3 | 57.7 | 44.5 | 16.0 | 70.4 | 66.8 |
| $C_2$ | 56.7 | 31.4 | 78.6 | 91.8 | 34.5 | 34.7 |
| $C_3$ | 42.3 | 53.8 | 84.8 | 99.2 | 79.0 | 44.6 |
| $C_4$ | 95.3 | 91.4 | 73.1 | 42.9 | 98.8 | 66.7 |
| $C_5$ | 76.2 | 98.4 | 31.0 | 55.4 | 14.5 | 57.4 |
| $C_6$ | 37.6 | 79.3 | 58.4 | 71.8 | 26.0 | 84.5 |

**TABLE V.** 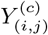 values of the 36 species for the 6C×6N×36S model.

| | $N_1$ | $N_2$ | $N_3$ | $N_4$ | $N_5$ | $N_6$ |
| --- | --- | --- | --- | --- | --- | --- |
| $C_1$ | 0.72 | 0.59 | 0.15 | 0.11 | 0.13 | 0.75 |
| $C_2$ | 0.29 | 0.72 | 0.79 | 0.39 | 0.15 | 0.16 |
| $C_3$ | 0.76 | 0.61 | 0.40 | 0.86 | 0.60 | 0.63 |
| $C_4$ | 0.87 | 0.68 | 0.64 | 0.27 | 0.80 | 0.51 |
| $C_5$ | 0.88 | 0.13 | 0.31 | 0.36 | 0.11 | 0.48 |
| $C_6$ | 0.46 | 0.34 | 0.38 | 0.83 | 0.70 | 0.86 |

**TABLE VI.** 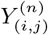 values of the 36 species for the 6C×6N×36S model.

| | $N_1$ | $N_2$ | $N_3$ | $N_4$ | $N_5$ | $N_6$ |
| --- | --- | --- | --- | --- | --- | --- |
| $C_1$ | 0.10 | 0.30 | 0.67 | 0.36 | 0.32 | 0.66 |
| $C_2$ | 0.83 | 0.30 | 0.47 | 0.30 | 0.40 | 0.58 |
| $C_3$ | 0.72 | 0.15 | 0.65 | 0.54 | 0.21 | 0.18 |
| $C_4$ | 0.30 | 0.22 | 0.84 | 0.64 | 0.29 | 0.56 |
| $C_5$ | 0.55 | 0.16 | 0.77 | 0.42 | 0.22 | 0.25 |
| $C_6$ | 0.49 | 0.67 | 0.89 | 0.80 | 0.50 | 0.19 |

## References

1 J. Qin, R. Li, J. Raes, M. Arumugam, K. S. Burgdorf, C. Manichanh, T. Nielsen, N. Pons, F. Levenez, T. Yamada, et al., nature 464, 59 (2010).

2 H. M. P. Consortium et al., Nature 486, 207 (2012).

3 J. M. Chaparro, A. M. Sheflin, D. K. Manter, and J. M. Vivanco, Biology and Fertility of Soils 48, 489 (2012).

4 Y. Pii, T. Mimmo, N. Tomasi, R. Terzano, S. Cesco, and C. Crecchio, Biology and Fertility of Soils 51, 403 (2015).

5 P. N. Hobson and C. S. Stewart, The rumen microbial ecosystem (Springer Science & Business Media, 2012).

6 R. D. Bardgett, C. Freeman, and N. J. Ostle, The ISME journal 2, 805 (2008).

7 A. Briones and L. Raskin, Current Opinion in Biotechnology 14, 270 (2003).

8 M. Wagner, A. Loy, R. Nogueira, U. Purkhold, N. Lee, and H. Daims, Antonie Van Leeuwenhoek 81, 665 (2002).

9 X. Zhou, C. J. Brown, Z. Abdo, C. C. Davis, M. A. Hansmann, P. Joyce, J. A. Foster, and L. J. Forney, The ISME journal 1, 121 (2007).

10 L. Lahti, J. Salojrvi, A. Salonen, M. Scheffer, and W. M. d. Vos, Nature Communications 5, ncomms5344 (2014).

11 C. A. Lozupone, J. I. Stombaugh, J. I. Gordon, J. K. Jansson, and R. Knight, Nature 489, 220 (2012).

12 J. Zhou, W. Liu, Y. Deng, Y.-H. Jiang, K. Xue, Z. He, J. D. V. Nostrand, L. Wu, Y. Yang, and A. Wang, mBio 4, e00584 (2013).

13 J. P. Sutherland, The American Naturalist 108, 859 (1974).

14 C. S. Holling, Annual review of ecology and systematics 4, 1 (1973).

15 R. M. May, Nature 269, 471 (1977).

16 T. Fukami and M. Nakajima, Ecology letters 14, 973 (2011).

17 T. Bush, M. Diao, R. J. Allen, R. Sinnige, G. Muyzer, and J. Huisman, Nature Communications 8, 789 (2017).

18 A. Schröoder, L. Persson, and A. M. De Roos, Oikos 110, 3 (2005).

19 A. Goyal, V. Dubinkina, and S. Maslov, The ISME journal, https://doi.org/10.1038/s41396 (2018).

20 D. Gonze, L. Lahti, J. Raes, and K. Faust, The ISME journal 11, 2159 (2017).

21 A. Shade, H. Peter, S. D. Allison, D. Baho, M. Berga, H. Buürgmann, D. H. Huber, S. Langenheder, J. T. Lennon, J. B. Martiny, et al., Frontiers in microbiology 3, 417 (2012).

22 T. J. Browning, E. P. Achterberg, I. Rapp, A. Engel, E. M. Bertrand, A. Tagliabue, and C. M. Moore, Nature 551, nature24063 (2017).

23 R. M. May, Nature 238, 413 (1972).

24 J. Mounier, C. Monnet, T. Vallaeys, R. Arditi, A.-S. Sarthou, A. Hélias, and F. Irlinger, Applied and environmental microbiology 74, 172 (2008).

25 S. Allesina and S. Tang, Nature 483, 205 (2012).

26 K. Faust and J. Raes, Nature Reviews Microbiology 10, 538 (2012).

27 R. R. Stein, V. Bucci, N. C. Toussaint, C. G. Buffie, G. Rätsch, E. G. Pamer, C. Sander, and J. B. Xavier, PLoS computational biology 9, e1003388 (2013).

28 S. Marino, N. T. Baxter, G. B. Huffnagle, J. F. Petrosino, and P. D. Schloss, Proceedings of the National Academy of Sciences 111, 439 (2014).

29 C. K. Fisher and P. Mehta, PloS one 9, e102451 (2014).

30 D. Berry and S. Widder, Frontiers in microbiology 5, 219 (2014).

31 K. Z. Coyte, J. Schluter, and K. R. Foster, Science 350, 663 (2015).

32 T. E. Gibson, A. Bashan, H.-T. Cao, S. T. Weiss, and Y.-Y. Liu, PLoS computational biology 12, e1004688 (2016).

33 J. Friedman, L. M. Higgins, and J. Gore, Nature ecology & evolution 1, 0109 (2017).

34 G. Bunin, Physical Review E 95, 042414 (2017).

35 Y. Fried, N. M. Shnerb, and D. A. Kessler, Physical Review E 96(2017), 10.1103/physreve.96.012412.

36 Y. Xiao, M. T. Angulo, J. Friedman, M. K. Waldor, S. T. Weiss, and Y.-Y. Liu, Nature communications 8, 2042 (2017).

37 A. J. Lotka, Elements of Physical Biology, by Alfred J. Lotka (1925).

38 V. Volterra, Animal ecology, 409 (1926).

39 B. Momeni, L. Xie, and W. Shou, Elife 6(2017).

40 R. MacArthur and R. Levins, Proceedings of the National Academy of Sciences 51, 1207 (1964).

41 R. MacArthur, Theoretical population biology 1, 1 (1970).

42 J. Huisman and F. J. Weissing, Ecology 82, 2682 (2001).

43 M. Tikhonov and R. Monasson, Physical Review Letters 118, 048103 (2017).

44 A. Posfai, T. Taillefumier, and N. S. Wingreen, Physical review letters 118, 028103 (2017).

45 J. E. Goldford, N. Lu, D. Bajić, S. Estrela, M. Tikhonov, A. Sanchez-Gorostiaga, D. Segrè, P. Mehta, and A. Sanchez, Science 361, 469 (2018).

46 S. Butler and J. O’Dwyer, bioRxiv, 293605 (2018).

47 D. Tilman, Monographs in population biology 17, 1 (1982).

48 A. Burson, M. Stomp, E. Greenwell, J. Grosse, and J. Huisman, Ecology 99, 1108 (2018).

49 D. N. Menge, L. O. Hedin, and S. W. Pacala, PLoS One 7, e42045 (2012).

50 V. S. Brauer, M. Stomp, and J. Huisman, The American Naturalist 179, 721 (2012).

51 T. Koffel, S. Boudsocq, N. Loeuille, and T. Daufresne, Ecology letters 21, 1010 (2018).

52 D. Gale and L. S. Shapley, The American Mathematical Monthly 69, 9 (1962).

53 D. Gusfield and R. W. Irving, The stable marriage problem: structure and algorithms (MIT press, 1989).

54 H. De Baar, Progress in Oceanography 33, 347 (1994).

55 G. F. Gause, Journal of experimental biology 9, 389 (1932).

56 G. F. Gauze, The struggle for existence (Baltimore, The Williams and Wilkins company, 1934).

57 A. Goyal and S. Maslov, Physical Review Letters 120, 158102 (2018).

58 Y. Fried, D. A. Kessler, and N. M. Shnerb, Scientific reports 6, 35648 (2016).

59 J. Grilli, M. Adorisio, S. Suweis, G. Barabás, J. R. Banavar, S. Allesina, and A. Maritan, Nature communications 8, 14389 (2017).

60 R. P. Rohr, S. Saavedra, and J. Bascompte, Science 345, 1253497 (2014).

61 S. S. Maslov, Unpublished (2018).

62 J. Milnor, Morse Theory, Vol. 51 (Princeton university press, 1963).

63 T. J. Case and R. G. Casten, The American Naturalist 113, 705 (1979).

64 P. Chesson, Theoretical Population Biology 37, 26 (1990).

64 V. D. Blondel, J.-L. Guillaume, R. Lambiotte, and E. Lefebvre, Journal of statistical mechanics: theory and experiment 2008, P10008 (2008).

66 A. C. Hindmarsh, P. N. Brown, K. E. Grant, S. L. Lee, R. Serban, D. E. Shumaker, and C. S. Woodward, ACM Transactions on Mathematical Software (TOMS) 31, 363 (2005).

67 K. Kovarova-Kovar and T. Egli, Microbiology and molecular biology reviews: MMBR 62, 646 (1998).

68 F. G. Bader, Biotechnology and Bioengineering 20, 183 (1978).

69 A. V. Tkachenko and S. Maslov, bioRxiv, 204826 (2017).

